# Impact of maternal compensation on developmental phenotypes in a zebrafish model of severe congenital muscular dystrophy

**DOI:** 10.1101/2025.05.13.653769

**Authors:** Kyle P. Flannery, Shorbon Mowla, Namarata Battula, L. Rose Clark, Deze Liu, Callista D. Oliveira, Cynthia Venkatesan, Lillian M. Simhon, Brittany F. Karas, Kristin R. Terez, Daniel Burbano-Lombana, M. Chiara Manzini

## Abstract

Genetic compensation is a common phenomenon in zebrafish in response to genetic alterations. As such, differences between morphant and mutant zebrafish models of human diseases have led to significant difficulties in phenotypic interpretation and translatability. One form of compensation is the maternal deposit of mRNAs and proteins into the oocyte that supports developmental processes before zygotic genome activation. In this study, we generated a zebrafish model of severe congenital muscular dystrophy by targeting *protein O-mannose N-Acetylglucosaminyltransferase 2* (*pomgnt2*), a maternally provided gene that maintains cell-extracellular matrix interactions through glycosylation. Zygotic knockouts (ZKOs) retain protein function in the first week post-fertilization and survive to adulthood, though they develop muscle disease later in life. In contrast, maternal-zygotic KOs (MZKOs) generated from ZKO females develop early-onset muscle disease, reduced motor function, neuronal axon guidance deficits, and retinal synapse disruptions, recapitulating features of the human presentation. While assessing transcriptional changes linked to disease progression, the availability of embryos obtained from different breeding strategies also allowed for direct comparison of ZKOs and MZKOs to define the impact of having a KO mother. We found that offspring from a ZKO mother, independently of genotype, show distinct expression patterns from animals obtained from heterozygous breeding. Some of these changes reflect an increased metabolic requirement, possibly stemming from maternal metabolic disruption. These findings will not only be applicable to other CMD models targeting maternally provided genes but also provide new insight into modeling disease using maternal-zygotic mutants.

## INTRODUCTION

Genetic compensation in zebrafish is a well-documented phenomenon that has profound implications for the use of zebrafish as a model system in studying developmental processes and human diseases [1]. The basis for genetic compensation arises from many sources, such as genome duplication in a teleost ancestor that led to additional copies of genes that are critical for embryonic development, as well as altered expression of non-duplicated, yet functionally similar genes that can mask the effect of genetic alterations [2–5]. As such, there has been considerable discrepancy between the phenotypes observed in fish following morpholino oligonucleotide (MO)-mediated knockdown strategies (morphants) and stable knock-out (KO) strains, leading to complications in phenotype interpretations [1, 6–9].

One such source of compensation in zebrafish stems from the maternal lineage, as female zebrafish provide an abundance of lipids, proteins, and other nutrients to the yolk of their externally fertilized oocytes that support the embryo and larva within the first 5 days post-fertilization (dpf) [10]. In addition to these nutrients, there are many maternally provided transcripts that direct embryonic development through the maternal to zygotic transition (MZT) when the zygotic genome is activated [11, 12]. In mutant models of human diseases, such as CHARGE syndrome and scoliosis, maternal transcripts have been found to compensate for loss of zygotic genes requiring the use of KO mothers to deplete maternal mRNAs for full phenotypic presentation [13, 14]. In the present study, we present a zebrafish KO strain for *protein O-linked mannose N-acetylglucosaminyltransferase 2* (*pomgnt2*) as a novel *in vivo* model of congenital neuromuscular disease also defining the impact of maternal compensation and declining maternal health in KO mothers.

*POMGNT2* encodes for a glycosyltransferase enzyme involved in the glycosylation of α-dystroglycan (α-DG), a ubiquitously expressed glycoprotein and the extracellular component of the dystrophin-glycoprotein complex (DGC) [15, 16]. α-DG links dystrophin and the intracellular cytoskeleton to ligands in the extracellular matrix (ECM) via an elongated functional glycan, termed matriglycan [17, 18]. Matriglycan is initiated via an O-linked mannose (O-Man) on the mucin domain of α-DG by the Protein O-mannosyltransferase 1 and 2 (POMT1, POMT2) complex and then extended by seven glycosyltransferases that add different sugars in a specific order. POMGNT2 catalyzes the addition of an N-acetylglucosamine to the O-Man and is selective for matriglycan over other O-Man initiated chains on α-DG [19, 20]. Biallelic variants in any of these glycosyltransferases cause a heterogenous group of congenital muscular dystrophies, termed dystroglycanopathies. Loss of function (LOF) variants in either *POMT1-2* or *POMGNT2* cause Walker Warburg Syndrome (WWS), the most severe dystroglycanopathy also associated with profound eye and brain malformations [21–23].

Due to the essential role of α-DG in placental formation in rodents, mouse KO models of multiple genes involved in α-DG glycosylation lead to embryonic or perinatal lethality, hindering studies of disease progression beyond birth [24–30]. As such, zebrafish have emerged as a prominent model of dystroglycanopathy. However, zebrafish KOs disrupting α-DG glycosylation present the same discrepancy between morphant and KO models that are prevalent in the field. Mutants for *dystroglycan* (*dag1*) itself, also known as *patchytail,* and another glycosyltransferase involved in matriglycan assembly, *fukutin-related protein* (*fkrp*), showed muscle disease and lethality within 10 dpf [31, 32]. In contrast, KOs for *pomt2* and *protein O-linked mannose N-acetylglucosaminyltransferase 1* (*pomgnt1*) have delayed onset months post fertilization [33, 34]. A recent study by our group reconciled these differences by showing the compensatory effects of maternal transcripts on the phenotypic presentation in a model for *POMT1* loss of function, indicating that the glycosyltransferases involved in α-DG glycosylation are maternally provided, and that α-DG glycosylation by maternally provided Pomt1 masks early developmental phenotypes [35]. We found the same to be true in developing a model of *POMGNT2* loss of function.

While generating maternal-zygotic mutants through zygotic mutant females has been widely undertaken to study the effect of maternally provided gene loss, there has not been as much investigation into the effects on offspring outcomes that may arise due to the zygotic mutations in female parents. However, when performing transcriptomic analyses to investigate disease progression, we also defined the differences between zygotic *pomgnt2* KO (ZKO) embryos that initially retain α-DG glycosylation and maternal-zygotic KOs (MZKOs) that do not. We revealed distinct correlations in gene expression patterns reflecting increased metabolic demand in MZKOs as well as their heterozygous siblings. These findings demonstrate for the first time how muscle wasting and declining health in zygotic mutant females may lead to physiological changes in their progeny and will have immediate applicability for the generation of novel zebrafish models of CMD and other genetic models that are impacted by maternal compensation.

## RESULTS

### Loss of zygotic pomgnt2 leads to adult-onset muscle phenotypes

To generate a *pomgnt2* KO strain, we used CRISPR-Cas9-induced nonhomologous end joining (NHEJ) with guide RNAs (gRNAs) targeting each of the two coding exons. We identified multiple frameshifts that disrupted *pomgnt2* in F1 founders. Deletions near the protospacer adjacent motif (PAM) sequence in exon 1 led to the usage of an alternative start codon, but we identified a 13 bp insertion (NM_001012384: c.17_29del, p.Cys6fs) with an additional 4 bp deletion (NM_001012384: c.32_35del, p.Pro11fs) that also disrupted the alternative start site. In addition, we found a 7 bp deletion in exon 2 in the glycosyltransferase domain (NM_001012384: c.713_719del, p.Ser238fs), and founders carrying both variants on the same allele (**Fig. 1A**). There are no suitable antibodies to test for Pomgnt2 protein expression in zebrafish. We tested KO embryos from heterozygous crosses (HetxHet) for nonsense mediated mRNA decay via qPCR and found that the mutated mRNA was present (**Supplemental Fig. 1A**). However, loss of Pomgnt2 function will lead to loss of matriglycan on α-DG. We confirmed by western blotting using the α-DG glyco-specific antibody clone IIH6C4 on glycoprotein-enriched lysates that α-DG glycosylation was absent in all three KO lines at 1 month post-fertilization (mpf), even when KO lysates were loaded in excess (**Fig. 1B**, **Supplemental Fig. 1B-C**). We used the line disrupting both exon 1 and exon 2 for further study.

**Figure 1.**
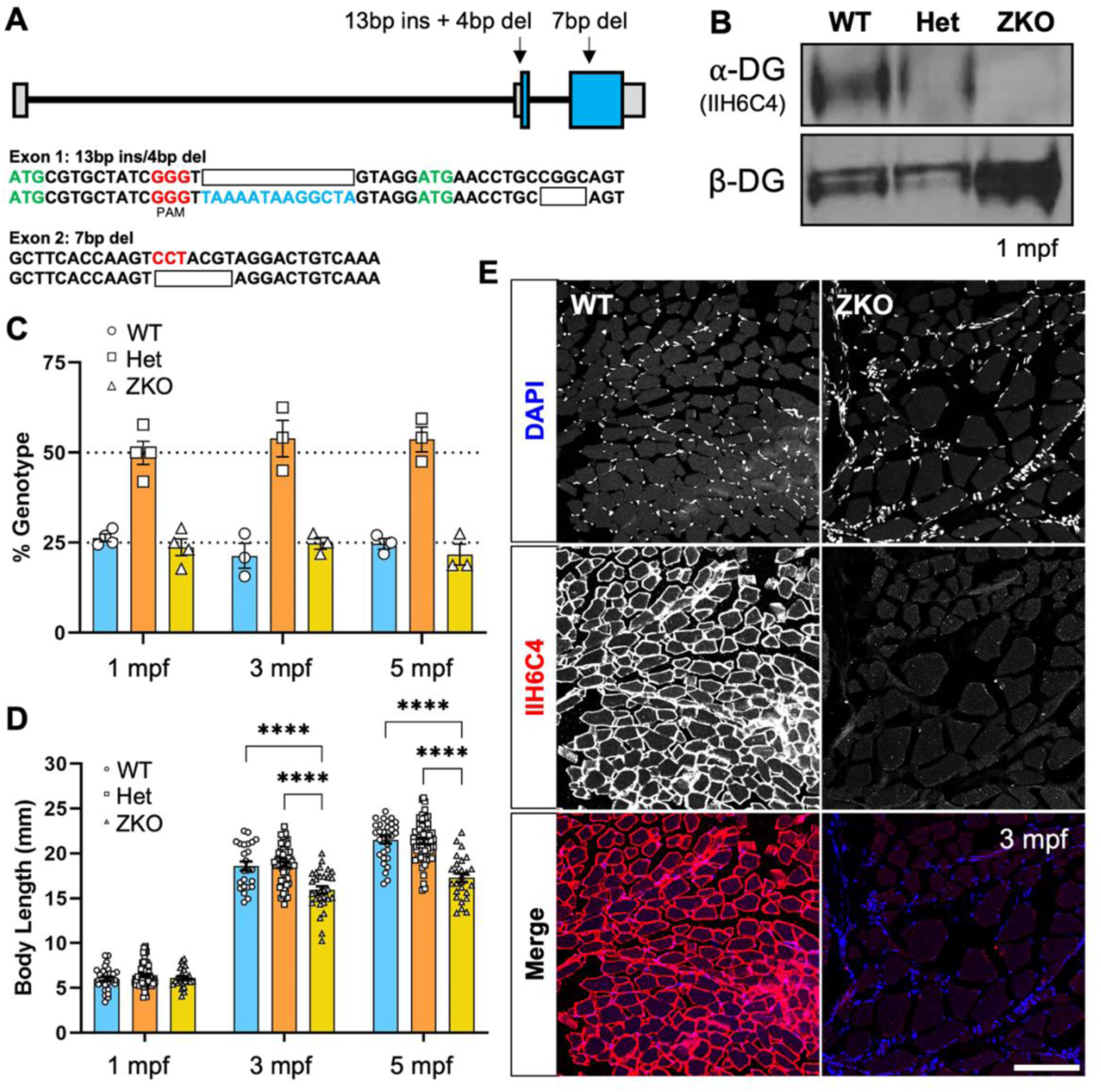
Generation of the pomgnt2 line and adult-onset phenotypes in ZKOs. **A**: Mutation schematic showing indels in exons 1 and 2 induced through CRISPR-Cas9 nonhomologous end joining. **B**: Western blot showing complete loss of α-DG glycosylation labeled via the glyco-specific IIH6C4 antibody in ZKOs at 1 mpf, even when glycoprotein-enriched lysate is in excess as shown through β-DG protein levels. **C**: Survival analysis showing ZKOs surviving in Mendelian ratios through 5 mpf. Expected survival rates of 25% for WTs and ZKOs and of 50% for Hets is indicated by dotted lines. **D**: Body length measurements showing comparable body length between ZKOs and their siblings at 1 mpf, but a significant reduction in ZKOs at 3 and 5 mpf. **E**: Fluorescent staining of transverse muscle sections showing loss of α-DG glycosylation (IIH6C4), fiber separation, and increased nuclear staining (DAPI) suggesting fibrosis and muscle disease in ZKOs (**Scale Bar**: 100 µm).

To determine how loss of *pomgnt2* impacts the overall health of the zebrafish, we examined survival, gross morphology, and muscle structure. We found that zygotic KOs (ZKOs) from heterozygous crosses (Het X Het) survive in Mendelian genotypic ratios into early adulthood (1 mpf: WT 26.35 ± 1.04%; Het 49.93 ± 3.23%; ZKO 23.72 ± 2.34 %; N=4 cohorts, n=134; 3 mpf: WT 21.32 ± 3.44%; Het 53.89 ± 5.05%; ZKO 24.79 ± 1.63%; N=3 cohorts, n=120; 5 mpf: WT 24.65 ± 1.51%; Het 53.68 ± 3.44%; ZKO 21.67±2.92 %; N=3 cohorts, n=120) (**Fig. 1C**). No differences in body length were observed at 1 mpf (**Fig. 1D**), but ZKOs were significantly smaller than their WT and Het siblings at 3 and 5 mpf (1 mpf: WT: 6.04±0.23 mm, n=30; Het: 6.41 mm ± 0.18, n=53; ZKO: 6.11 mm ± 0.20, n=26; N=3 cohorts. p>0.9999; 3 mpf: WT: 18.61 mm ± 0.51, n=24; Het: 18.76 mm ± 0.29, n=57; ZKO: 15.93 mm ± 0.41, n=30; ****p<0.0001. 5 mpf: WT: 21.53 mm ± 0.40, n=30; Het: 21.33 mm ± 0.28, n=62; ZKO: 17.29 mm ± 0.47, n=26; N=3 cohorts. ****p<0.0001) (**Fig 1D**).

One of the main features of dystroglycanopathy is loss of muscle integrity which mirrored the delayed growth. No overt muscle phenotype was found at 1 mpf. Muscle fibers stained with fluorescently conjugated phalloidin were organized and uniform in ZKOs, comparable to WT (**Supplemental Fig. 1D**). General signs of muscle disease were present at 3 mpf with hallmark features such as variation in myofiber size, gaps between myofibers, and increased nuclear staining suggesting fibrosis (**Fig. 1E**). While survival was only formally assessed through 5 mpf, ZKOs could be maintained up to approximately 1 year of age at lower stocking densities to limit food competition. At this time, muscle disease was significantly advanced with evidence of severe fibrosis (**Supplemental Fig. 2A**), and greatly disrupted locomotor function in ZKOs (**Supplemental Fig. 2B-D**). Taken together, these findings indicate that ZKOs exhibit prolonged survival but have progressive muscle disease in adulthood.

### Loss of maternal and zygotic pomgnt2 unmasks early developmental phenotypes

While loss of α-DG itself in *dag1* ZKOs leads to severe phenotypes during the first two weeks post fertilization [36], the mild phenotypic presentation in *pomgnt2* ZKOs mirrored previous findings in *pomt1* ZKOs undergoing maternal compensation [36]. *pomgnt2* mRNA has previously been detected in the embryo at developmental stages before the MZT indicating that it is maternally provided [37]. We confirmed that *pomgnt2* ZKOs retain α-DG glycosylation in embryos and larvae by examining IIH6C4 immunostaining at 7 dpf (**Fig. 2A**). Therefore, we bred ZKO females with Het males resulting in progeny with an expected ratio of 50% maternal *pomgnt2* Hets (MHets) and 50% maternal zygotic KOs (MZKOs). Here, we found that α-DG glycosylation was eliminated in MZKOs at 7 dpf (**Fig. 2A**), supporting our hypothesis that maternal compensation was the most likely source of residual glycosylation in ZKOs.

**Figure 2.**
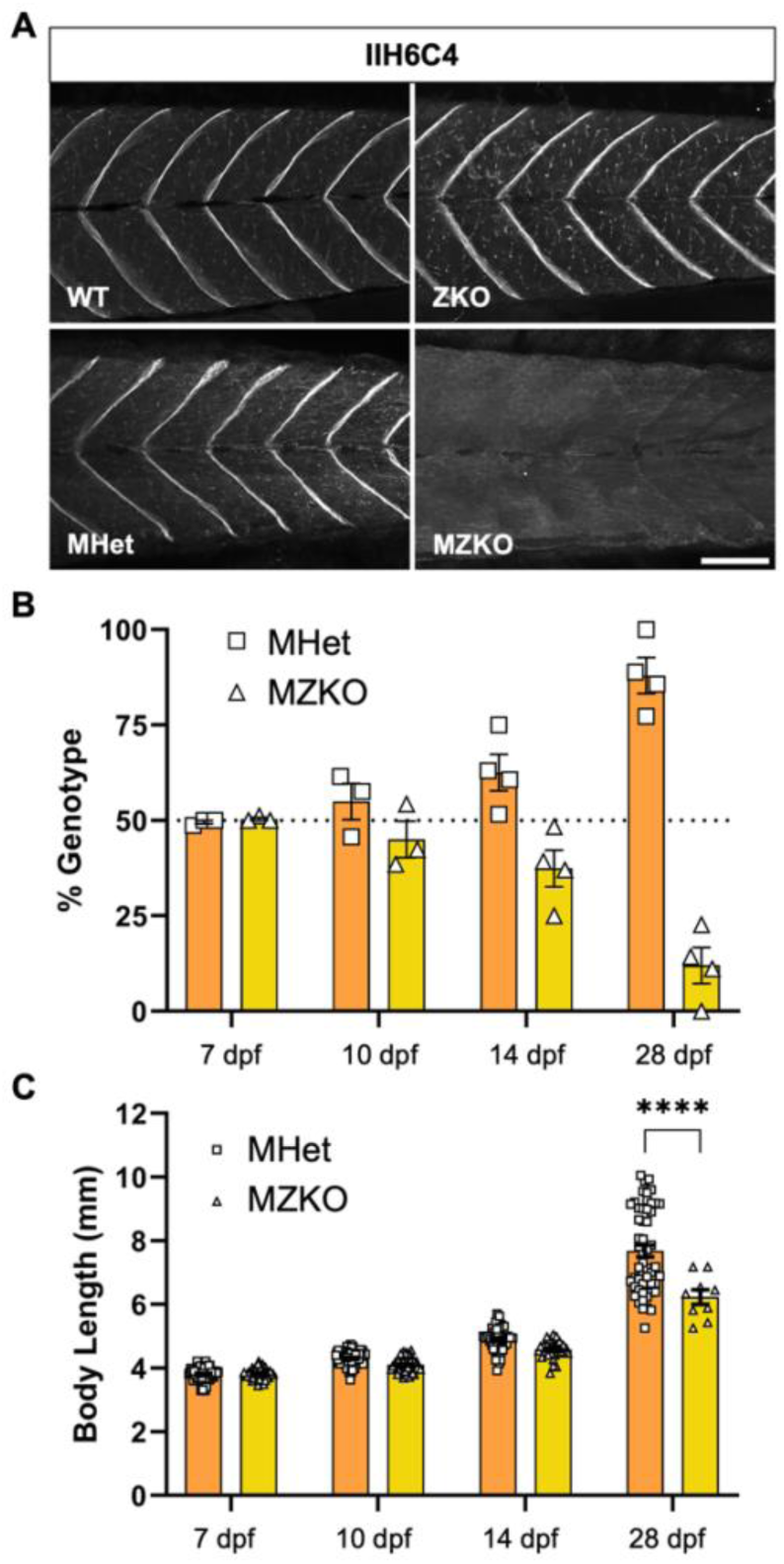
Developmental phenotypes caused by elimination of maternal pomgnt2. **A**: Immunofluorescent staining of muscle showing that ZKOs have residual α-DG glycosylation due to maternally provided *pomgnt2*, but this is depleted in MZKOs without maternal *pomgnt2* (**Scale Bar**: 100 µm). **B**: Survival analysis showing that MZKOs begin to deviate from the Mendelian 50/50 survival ratio between 10-14 dpf, with most MZKOs dead by 28 dpf. **C**: Body length measurements showing a trend toward reduction in MZKOs at 14 dpf and a significant reduction at 28 dpf.

As expected, MZKOs survival progressively declined between 10 and 14 dpf and the vast majority of MZKOs died by 28 dpf (7 dpf: MHet 49.57 ± 0.43%; MZKO 50.40 ± 0.40%; N=3 cohorts. 10 dpf: MHet 54.93 ± 4.75%; MZKO 45.07 ± 4.75%; N=3 clutches. 14 dpf: MHet 62.18 ± 5.13%, n=42; MZKO 37.83 ± 5.13%; N=4 clutches. 28 dpf: MHet = 87.98 ± 4.70%, MZKO = 12.03 ± 4.70%; n=61, N=4 clutches) (**Fig. 2C**). Data from each timepoint were examined collectively by Chi-square analysis, indicating a statistically significant deviation in survival beginning at 14 dpf that increases drastically by 28 dpf (14 dpf: MHet n=59, MZKO n=39, χ^2^=4.08, df=1, *p=0.04. 28 dpf: MHet n=52, MZKO n=9, χ^2^=29.4, df=1, ****p<0.0001) (**Supplemental Table 1**). Trends towards reduced body length were noted at 10 dpf and 14 dpf. The few MZKOs that survived to 28 dpf were significantly smaller than MHets (**Fig.2C**) (7 dpf: MHet 3.82 ± 0.03 mm, n=67; MZKO 3.83 ± 0.03 mm, n=42; p>0.9999. 10 dpf: MHet 4.32 mm ± 0.03, n=66; MZKO 4.11 ± 0.03 mm, n=55; N=3 clutches; p=0.1611. 14 dpf: MHet 4.90 ± 0.06 mm, n=42, MZKO 4.60 ± 0.06 mm, n=30; N=3 clutches; p=0.0857; 28 dpf: MHet 7.68 ± 0.19 mm, n=52; MZKO 6.24 ± 0.23 mm, n=9; N=4 clutches; ****p<0.0001). Collectively, these assessments show that removing maternal *pomgnt2* mRNA reveals developmental phenotypes in MZKO larvae.

### MZKOs show dystroglycanopathy phenotypes within the first 2 weeks post-fertilization

Severe dystroglycanopathy presents as loss of mobility and muscle integrity, retinal abnormalities, and neuronal axon guidance deficits in fish models [31, 32, 35]. To determine the onset of dystroglycanopathy phenotypes in MZKOs, we performed multiple muscle integrity and function analyses. Locomotor activity analysis using automated tracking in open swimming trials found that at 5 dpf, before the onset of any differences in body size, MZKOs already exhibited profound deficits in swimming behaviors. Significant differences were found in total swimming distance (**Fig. 3A**) (MHet: 375.2 ± 12.13 cm, n=79; *MZKO*: 228.4 ± 58.58 cm, n=91; ****p<0.0001), average velocity (**Fig. 3B**) (MHet: 0.21 ± 0.007 cm/s, n=79; MZKO: 0.13 ± 0.003 cm/s, n=90; ****p<0.0001), maximum velocity (**Supplemental Fig. 3A**) (MHet: 7.42 ± 0.24 cm/s, n=79; MZKO: 2.23 ± 0.24 cm/s, n=90; ***p<0.001), and maximum acceleration (**Supplemental Fig. 3B**) (MHet: 191.4 ± 7.07 cm/s^2^, n=79; MZKO: 155.6 ± 6.83 cm/s^2^, n=90; ***p<0.001).

**Figure 3.**
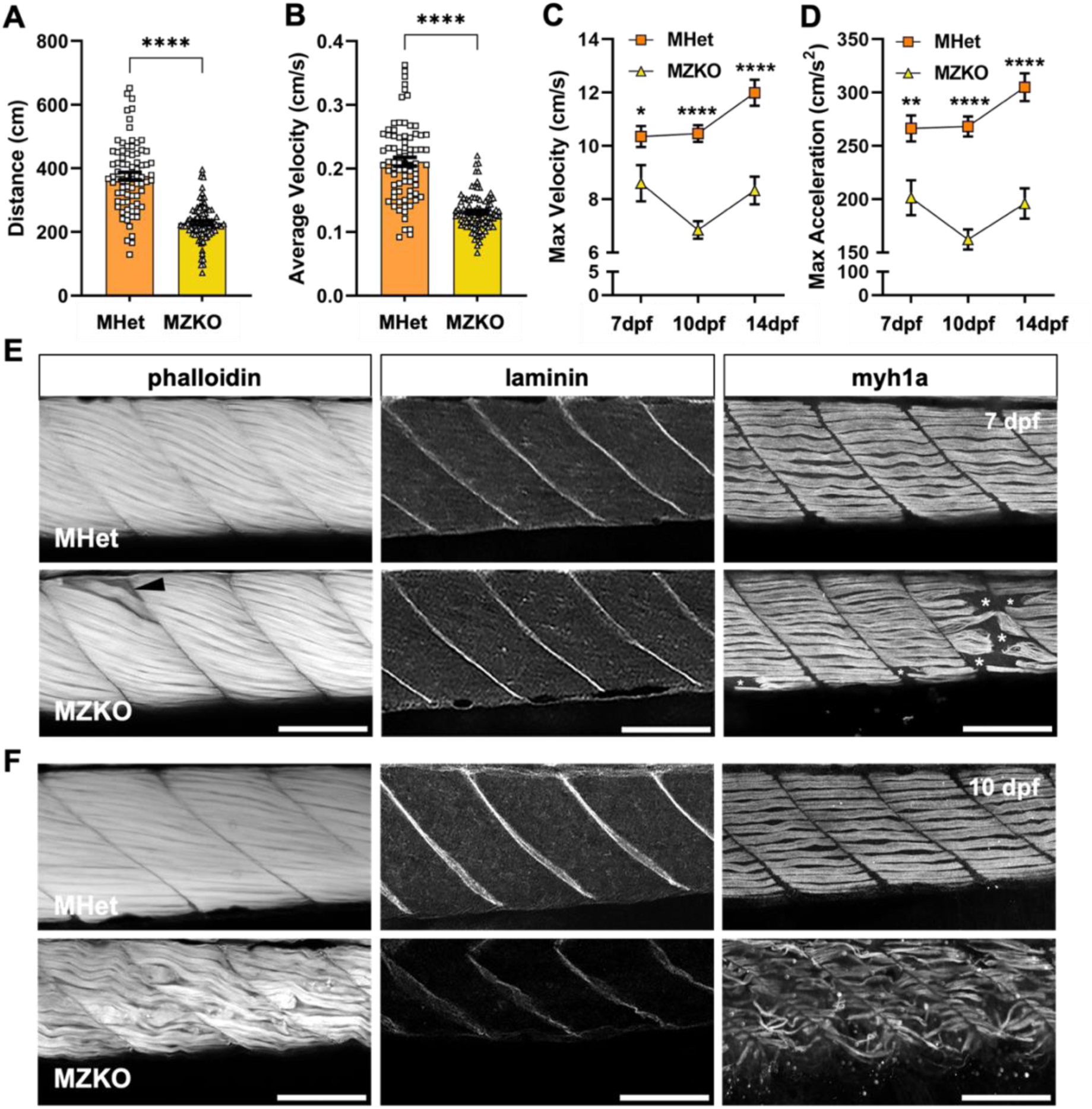
Early onset muscle and motor phenotypes in MZKOs. **A-B**: Analysis of locomotor function at 5 dpf showing significant reductions in distance (**A**) and average velocity (**B**) in MZKOs. **C-D**: Analysis of locomotor function between 7-14 dpf showing that MZKOs have significant reductions in maximum velocity (**C**) and acceleration (**D**) at all three timepoints compared to their MHet siblings but show an overall decline between 7-10 dpf. **E**: Fluorescent staining of muscle at 7 dpf showing overall uniform muscle organization with a few detached fibers (**left**, arrowhead), normal laminin signal at the myotendinous junctions (**middle**), and detachments within the slow twitch fiber layer (**right**, asterisks). **F**: Fluorescent staining of muscle at 10 dpf showing overall deterioration of muscle fibers (**left**), diffuse and disrupted laminin signal at the myotendinous junctions (**middle**), and uniform degeneration of the slow twitch fiber layer (**right**). **All Scale Bars**: 100 µm.

We further leveraged locomotor behavior as a readout of muscle function at 7, 10, and 14 dpf with a larger swimming arena to help define the course of disease progression. A significant reduction in distance and average velocity was noted at all three timepoints (**Supplemental Fig. 3C-D**), but there was an increasing difference in maximum velocity and maximum acceleration between MHets and MZKOs between 7 and 14 dpf suggesting that a drastic decline in muscle strength after 7 dpf (**Fig. 3C-D**).

Despite the presence of motor deficits at 7 dpf, integrity of actin filaments in muscle fibers labeled with fluorescently conjugated phalloidin (**Fig. 3E**) and of the laminin basement membrane at the myotendinous junctions (MTJs) (**Fig. 3F**) was generally unaffected in MZKOs, with the exception a few occasional detached myofibers (**Fig. 3E**). Interestingly, immunostaining for slow-twitch muscle fibers using myosin heavy chain 1A (Myh1a – F59 clone) revealed sparse detachment at the MTJ in some myotomes, similar to the early fiber detachment found in Duchenne Muscular Dystrophy (DMD) and congenital muscular dystrophy models (**Fig. 3E**) [36]. By 10 dpf, muscle disease in MZKOs had progressed rapidly, with several myotomes displaying detached, atrophied, and disorganized muscle fibers, disrupted and split MTJs, and complete deterioration of Myh1a-positive fibers (**Fig. 3F**).

We next examined the structure of the neuromuscular junctions (NMJs), as Agrin, a major contributor to acetylcholine receptor (AChR) clustering, is a well-characterized α-DG binding partner [38–40]. We visualized AChRs using fluorescently labeled α-bungarotoxin (α-BTX) and motor neuron terminals via immunostaining for synaptic vesicle protein 2 (SV2) at 10 dpf (**Fig. 4A**). No significant differences were noted between MHets and MZKOs in the myotome in α-BTX intensity or density and in SV2 puncta density (**Fig. 4B-D**) (α-BTX intensity: MHet: 0.1324 ± 0.010, n=13; MZKO: 0.1082 ± 0.007, n=9; p=0.0605. α-BTX puncta density: MHet: 1.43 ± 0.10 puncta/100 µm^2^, n=14; MZKO: 1.62 ± 0.21 puncta/100 µm^2^, n=10; p=0.3806. SV2 puncta density: MHet: 2.04 ± 0.09 puncta/100 µm^2^, n=14; MZKO: 1.91 ± 0.13 puncta/100 µm^2^, n=10; p=0.3490). However, there was a significant reduction in ⍺-BTX/SV2 colocalization, indicating a disruption in NMJ synapse integrity that could impact muscle innervation (MHet: 0.75 ± 0.01, n=14; MZKO: 0.6923 ± 0.01, n=10; ***p=0.004) (**Fig. 4E**). AchR cluster fragmentation was more evident at the MTJ, where we observed a significant reduction in α-BTX intensity paired with an increase in α-BTX puncta density (**Fig.4B’-C’**) (α-BTX intensity: MHet: 0.1548 ± 0.006, n=12; MZKO: 0.1083 ± 0.005, n=9; ****p=0.0001. α-BTX puncta density: MHet: 4.86 ±0.11 puncta/100 µm^2^, n=14; MZKO: 5.85 ± 0.13 puncta/100 µm^2^, n=10; ****p<0.0001). SV2 puncta density was not significantly different (**Fig.4D’**) (MHet: 3.92 ± 0.23 puncta/100 µm^2^, n=14; MZKO: 3.42 ± 0.17 puncta/100 µm^2^, n=10; p=0.1258). Lastly, α-BTX/SV2 colocalization showed a larger reduction at the MTJ than in the myofibers (**Fig. 4E’**) (MHet: 0.78 ± 0.01, n=14; MZKO: 0.6462 ± 0.01, n=10; ***p<0.0001). Taken together, these data indicate that loss of *pomgnt2* affects the NMJs, with AchR fragmentation at the MTJ in end plates primarily innervated by secondary motor neurons.

**Figure 4.**
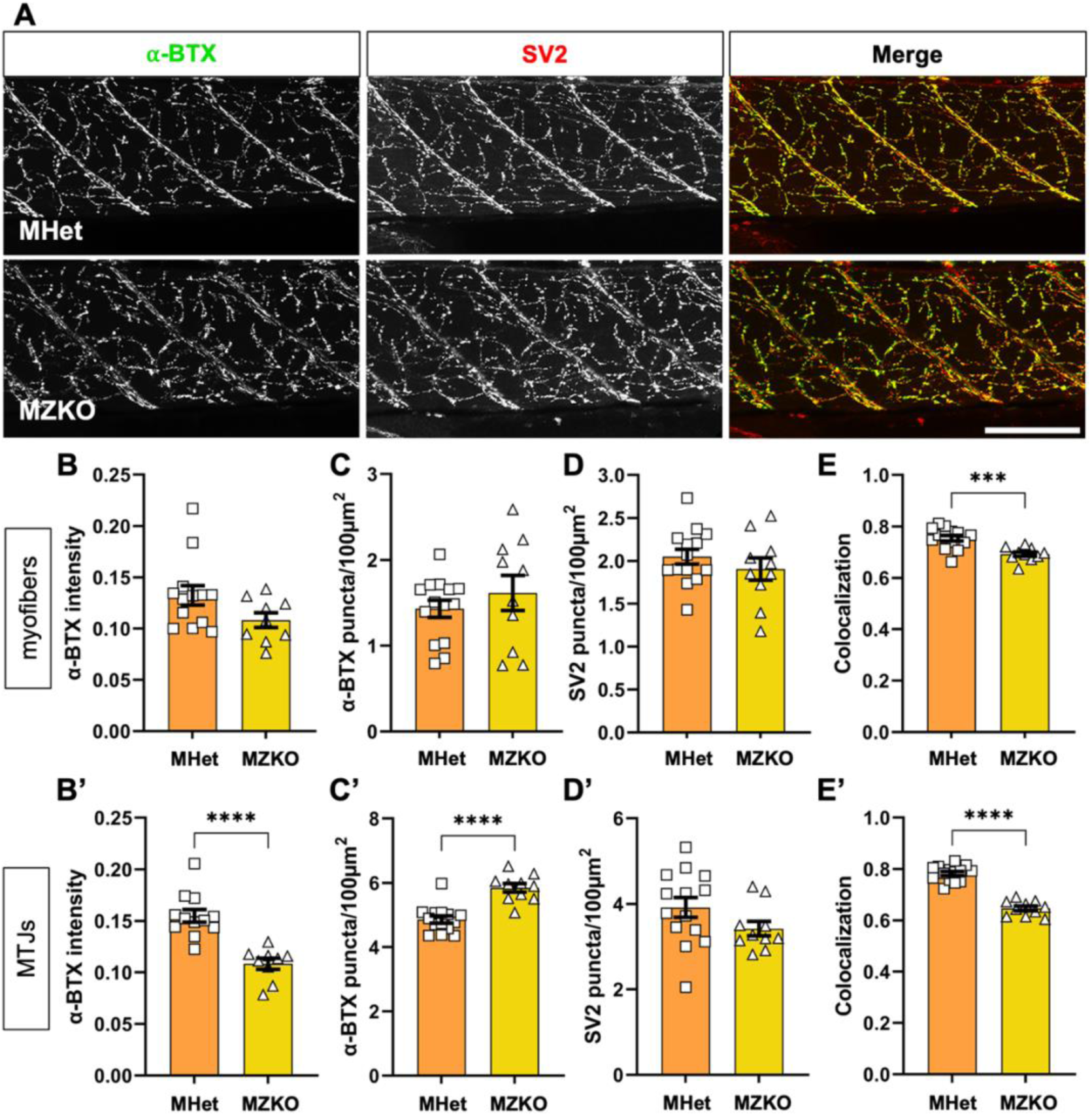
Evaluation of neuromuscular junction integrity in MZKOs. **A:** Fluorescent staining of muscle with α-bungarotoxin (BTX) to label Acetylcholine receptors on the sarcolemma (**left**), anti-SV2 antibody to label motor neuron terminals (**middle**), and merged images to evaluate colocalization (**right**) (**Scale Bar**: 100 µm). **B-D**: Quantifications of neuromuscular junctions within the myotomes showing no significant differences in α-BTX intensity (**B**), α-BTX puncta density (**C**), or SV2 puncta density (**D**), but a significant reduction in colocalization (**E**) in MZKOs. **B’-D’**: Quantifications of neuromuscular junctions at the myotendinous junctions showing significantly reduced α-BTX intensity (**B’**), significantly increased α-BTX puncta density (**C’**), no change in SV2 puncta density (**D’**), and a significant reduction in colocalization (**E’**).

Retinal photoreceptor synapse loss is another hallmark feature of dystroglycanopathies recapitulated in zebrafish models [33, 34]. We examined how the 10 dpf retina is impacted by immunostaining with an anti-Arrestin-3a (Arr3a) antibody (zpr1 clone) to outline both the outer segment and pedicles of cone photoreceptors, and with an anti-synaptophysin antibody to identify presynaptic vesicles at ribbon synapses in the photoreceptor pedicles. Synaptophysin staining showed occasional discontinuities and reduced intensity in the outer plexiform layer (OPL) (MHet: 0.1668 ± 0.016, n=9; MZKO: 0.0701 ± 0.021, n=7; **p=0.0022) (**Fig.5A-B**). In addition, photoreceptor cell bodies in the outer nuclear layer (ONL) were often less organized and occasionally protruded into the OPL, though the outer segment of the cones showed comparable organization between MHets and MZKOs indicating that these disruptions to the inner retinal layers have not yet led to photoreceptor death. Horizontal cell nuclei lining the upper border of the inner nuclear layer (INL) also appeared more diffuse and disorganized in MZKOs (**Fig. 5A**).

**Figure 5.**
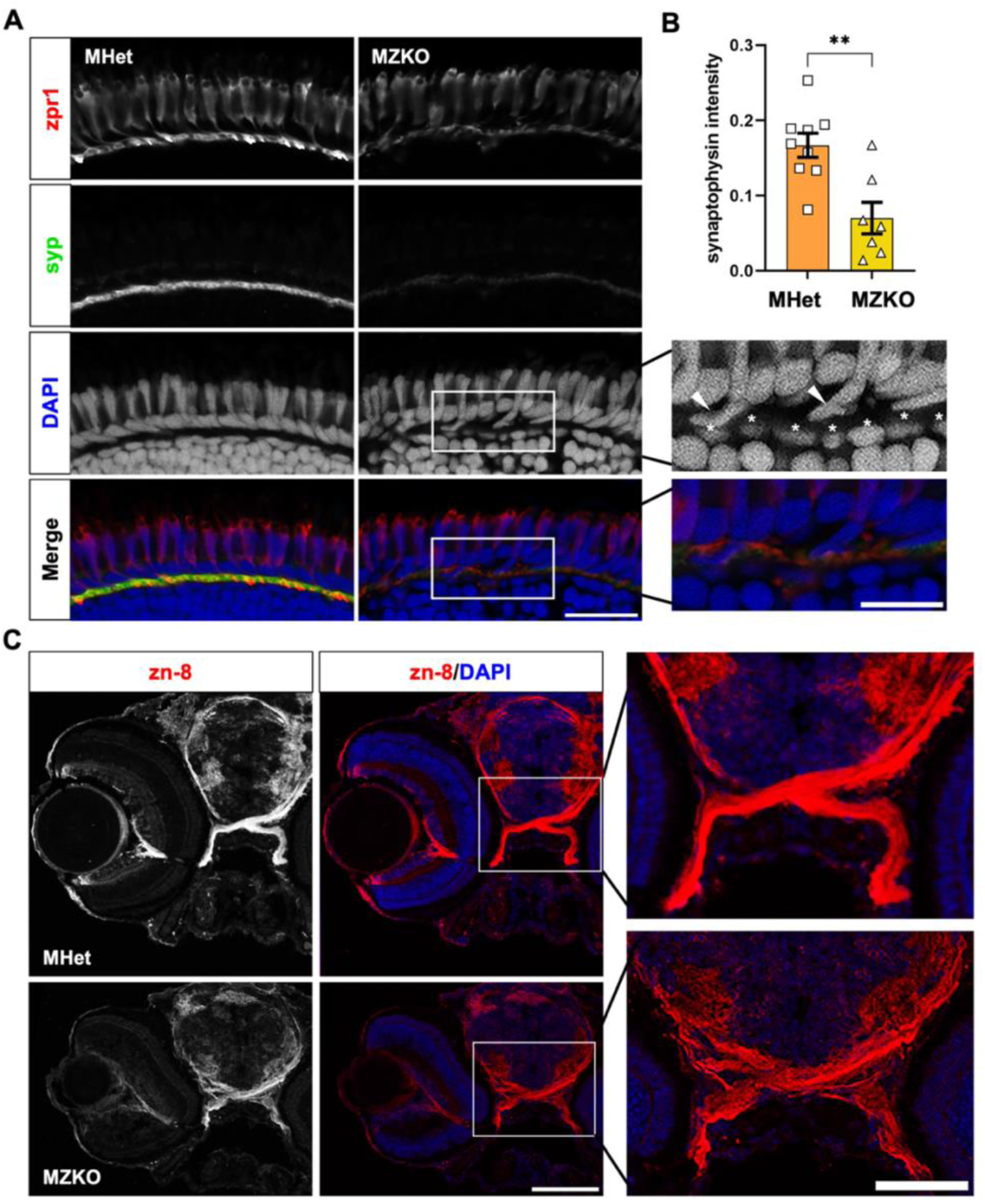
Retinal synapse formation and axon guidance deficits in MZKOs. **A**: Fluorescent staining of the outer layers of the retina showing comparable zpr1 (Arr3a) staining in the photoreceptor outer segments and pedicles in the outer plexiform layer, but reduced and disrupted synaptophysin (syp) staining and protrusion of photoreceptor cell bodies in the outer plexiform layer (arrowheads) and disorganization of horizontal cells lining the top of the outer nuclear layer (asterisks) suggesting defects to the ribbon synapses (**Scale Bars: 20 µm** for main panels, **10 µm** for zoomed in DAPI and merged MZKO panels). **B**: Quantification of synaptophysin intensity in the outer plexiform layer showing a significant reduction in MZKOs. **C**: Fluorescent staining of transverse cryosections with zn-8 and DAPI showing an overall disruption in morphology of the retina with evidence of retinal ganglion cell axon defasciculation in the optic nerves at the chiasm (**Scale Bars**: **100 µm** for images of whole retina, **20 µm** for zoomed in images of the chiasm).

Finally, we examined the structure of the optic chiasm, as α-DG plays a role in axon guidance through interactions with the Slit family of axon guidance cues that facilitate midline crossing [41, 42]. Used the zn-8 antibody to label retinal ganglion cell axons, we found evidence of defasciculation that was most prominent following decussation of the optic nerves, as was observed in *pomt1* and *slit2* KOs (**Fig. 5C**) [35, 43]. This suggests that loss of *pomgnt2* impacts the retina and axon guidance in a manner similar to other genes involved in α-DG glycosylation. Overall, MZKOs recapitulate features of dystroglycanopathy and closely resemble other severe CMD models.

### Loss of maternal pomgnt2 leads to differences in metabolic gene expression

To investigate both the molecular changes involved in disease progression in MZKOs and potential sources of compensation in ZKOs, we performed transcriptomic analyses at different ages. To define changes associated with the severe disease state when MZKOs begin to die, we conducted bulk RNA sequencing (RNA-seq) analyses on samples obtained from whole 10 dpf larvae, comparing them with MHet siblings. We identified 959 differentially expressed genes (DEGs), with 810 significantly downregulated and 149 significantly upregulated DEGs (padj<0.01; FC<0.7) (**Fig. 6A**).

**Figure 6.**
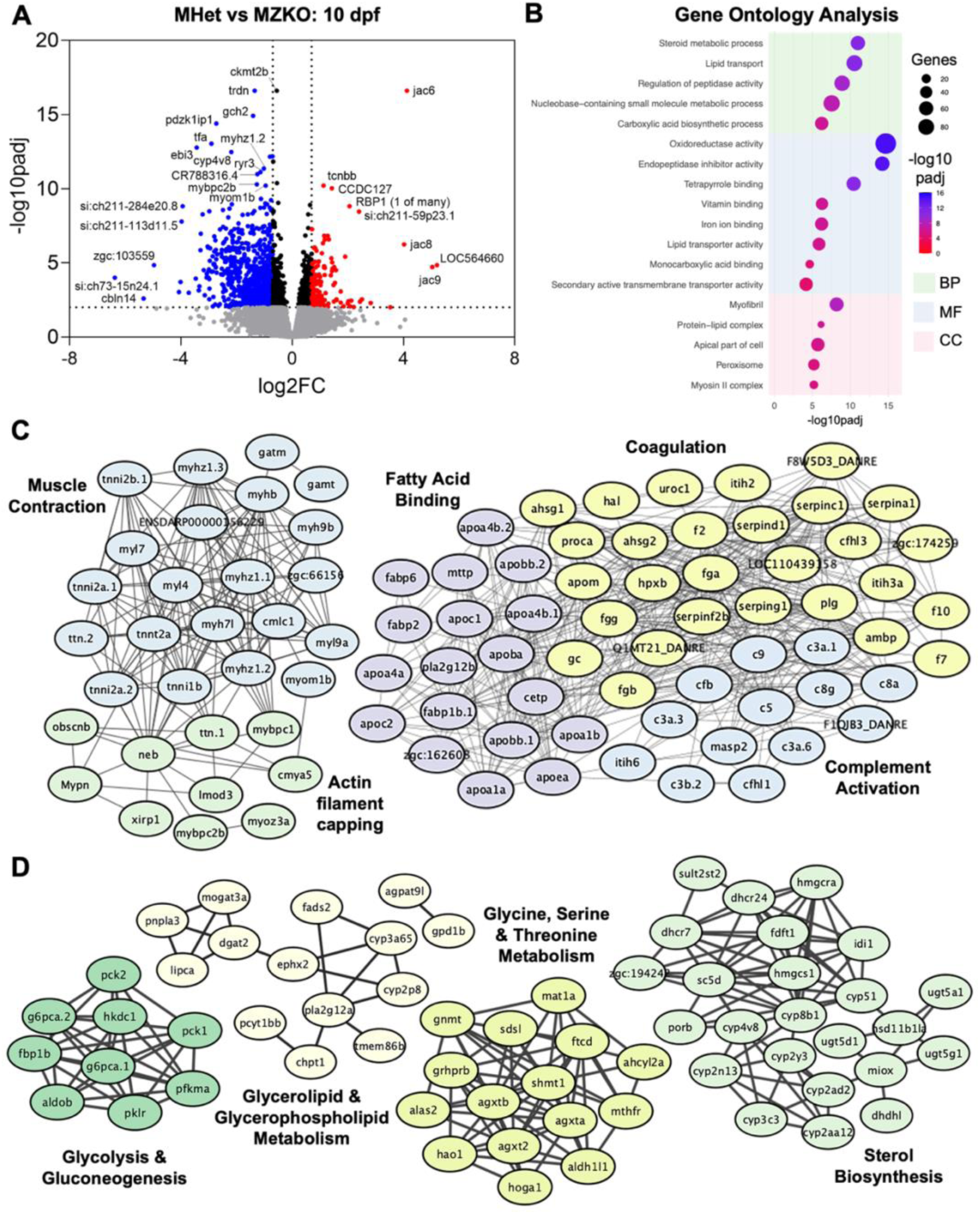
Differential gene expression analysis in MHets and MZKOs at 10 dpf. **A**: Volcano plot showing 810 significantly downregulated DEGs and 149 significantly upregulated DEGs. **B**: Gene Ontology analysis showing various enriched Biological Processes, Molecular Functions, and Cellular Components among the downregulated DEGs pertaining to lipid and cholesterol metabolism and muscle differentiation and function. **D-E**: STRING local networks analysis of interactions among the protein products of the downregulated DEGs showing a large, interconnected network of proteins involved in muscle contraction and actin filament capping, as well as fatty acid binding, coagulation, and complement activation (**D**), and smaller networks of proteins involved in glycolysis and gluconeogenesis, glycerophospholipid metabolism, amino acid metabolism (glycine, serine, threonine), and sterol biosynthesis (**E**).

Gene Ontology (GO) analysis showed a large number of enriched biological processes (BPs), many of which were redundant and required curation through analysis of overlapping genes. The most highly enriched BPs, molecular functions (MFs) and cellular components (CCs) were related to muscle function, metabolism, and regulation of proteolytic activity among the downregulated DEGs (**Fig. 6B**). Increased expression of *jacalin* family genes led to GO enrichment in mannose binding among the upregulated DEGs. This analysis was further refined using local networks defined by protein-protein interactions in STRING, which revealed several additional downregulated networks. These included lager interconnected networks including muscle contraction and actin filament capping, in addition to complement activation, coagulation, fatty acid binding, phospholipid metabolism, and LDL remodeling. The latter was driven by reduced expression of many apolipoprotein transcripts (*apoa1a, apoa1b, apoea, apoa4a, apoa4b.1, apoa4b.2, apoc2, apom*) (**Fig. 6C**). Additional distinct networks of downregulated genes included those involved in glycolysis and gluconeogenesis, glycerophospholipid metabolism, sterol/cholesterol biosynthesis, and amino acid (glycine, serine, threonine) metabolism (**Fig. 6D**). These findings suggest that MZKOs are in a severe state of muscle disease with global disruptions in metabolic processes.

MZKOs were also tested at an earlier timepoint at 5 dpf when mobility is reduced but muscle integrity is preserved and further compared with the progeny of Het x Het crosses. ZKOs still show WT-levels of glycosylation due to maternal compensation (**Fig. 2A**) and we wanted to test whether additional compensatory changes were present. Transcriptional dysregulation was much more modest in ZKOs, than MZKOs (**Fig. 7A** and **C**). However, ZKOs still showed 17 upregulated and 203 downregulated DEGs when compared to WTs (**Fig. 7A**). Among the upregulated genes were the transcription factors *junba, junbb, fosab,* and *egr2a*, which are involved in signaling cascades that promote cell proliferation and differentiation. The glucuronosyltransferase *b3gat1b* which is involved in the synthesis of a unique sulfated trisaccharide, the HNK-1 epitope, was also upregulated possibly indicating a compensatory response to increase cell-cell and cell-ECM interactions independent of α-DG [44]. Downregulated DEGs, belonged to networks involved in cation homeostasis, complement activation, and metabolism of carbon, amino acids, and phospholipids indicating some metabolic disruptions (**Fig.7B**). A small network of genes involved in the formation of the myosin II complex involved in were also noted among the downregulated DEGs (**Fig. 7B**).

**Figure 7.**
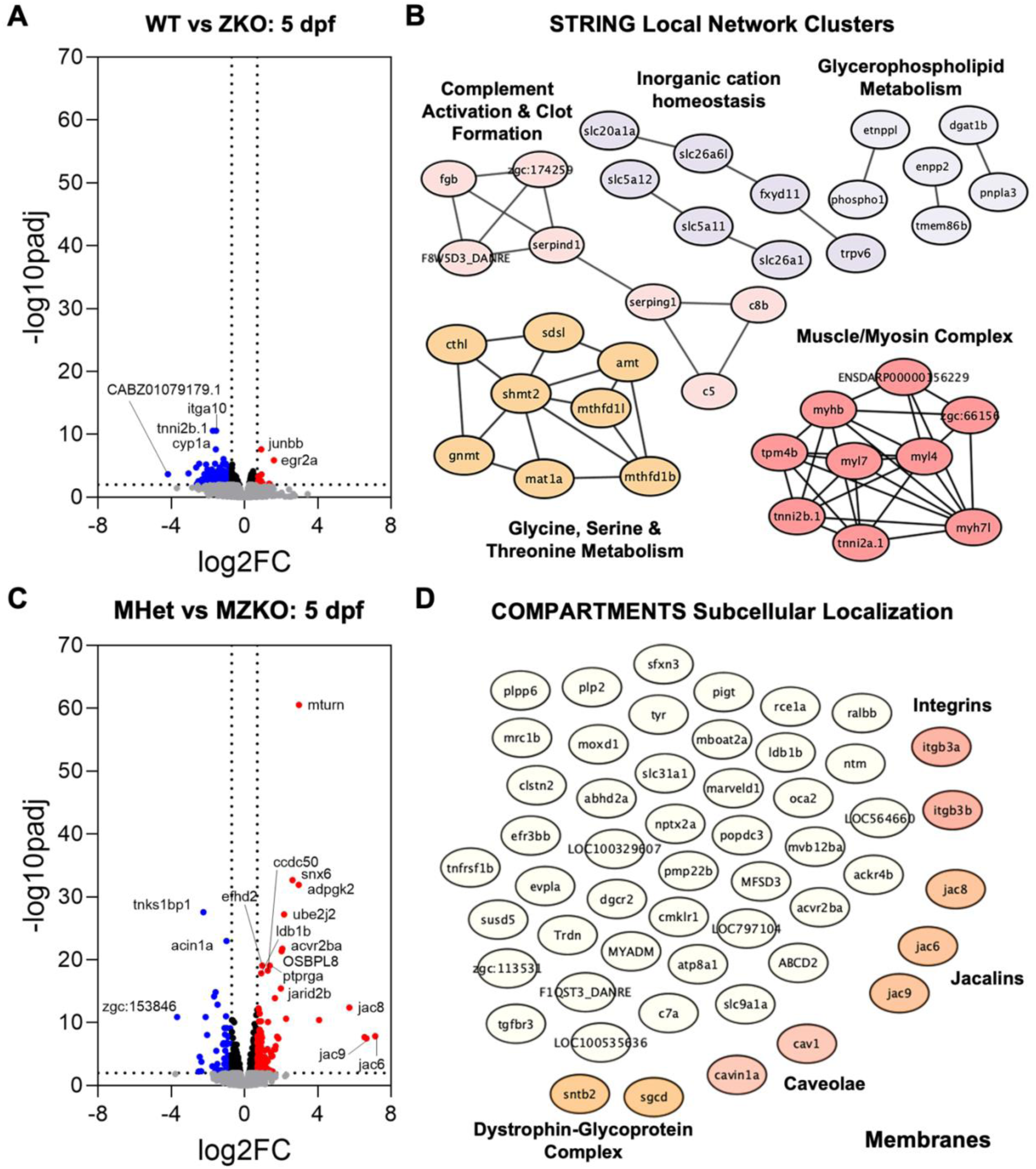
Differential gene expression analysis at 5 dpf. **A**: Volcano plot showing 203 downregulated DEGs and 17 upregulated DEGs in ZKOs compared to their WT siblings. **B**: STRING local networks analysis of downregulated DEGs showing networks of interacting proteins involved in complement activation and clot formation, inorganic cation homeostasis, glycerophospholipid metabolism, amino acid metabolism (glycine, serine, threonine), and muscle function within the myosin II complex. **C**: Volcano plot showing 84 downregulated DEGs and 159 upregulated DEGs in MZKOs compared to their MHet siblings. **D**: Subcellular localization analysis of the downregulated DEGs in **C** through COMPARTMENTS showing an enrichment in genes that encode proteins that are present in the membrane.

5 dpf MZKOs showed a distinct transcriptional disruption with 159 upregulated and 84 downregulated DEGs compared to MHets (**Fig. 7C**). While there was no enrichment for GO BPs, COMPARTMENTS analysis for subcellular localization along with several enriched GO CCs indicated that many upregulated DEGs are present on membranes. Plasma membrane proteins included *sarcoglycan δ* (*sgcd*) and *syntrophin β2* (*sntb2*), which promote membrane stability within the DGC, as well as several *integrins* (*itgb3a, itgb3b*) (**Fig. 7D**), suggesting a compensatory response to stabilize cell-ECM interactions independent from matriglycan. In parallel, negative regulators of muscle development such as *eif4ebp3I* and *ssh2a* were downregulated while *fbxo32*, which is linked to muscle wasting, was upregulated. Collectively, the transcriptomic analyses showed that MZKOs are in a severe state of muscle and metabolic disease at 10 dpf, and some disease-relevant pathways become dysregulated in both ZKOs and MZKOs before the full onset of disease phenotypes. Though, ZKOs and MZKOs display different patterns of transcriptional dysregulation that are consistent with preserved α-DG glycosylation in ZKOs.

To further probe whether the DEG expression patterns in the different genotypes at 5 dpf could reveal additional biological changes, we used weighted gene co-expression network analysis (WGCNA) comparing the three genotypes obtained from HetxHet crosses, WT, Het and ZKO, and from KOxHet crosses, MHet and MZKO. Following dataset reduction and soft thresholding (**Supplemental Fig. 4A-B**), this analysis identified seven modules comprised of 1818 genes total whose expression was strongly correlated (**Supplemental Fig. 4C-D**). Four of the seven gene modules were strongly correlated with specific genotypes and crosses (**Fig. 8A**), with a positive correlation indicating increased expression and a negative correlation indicating decreased expression (**Supplemental Fig. 4E-H**). Surprisingly, we identified modules that were driven by either disease state or maternal state.

**Figure 8.**
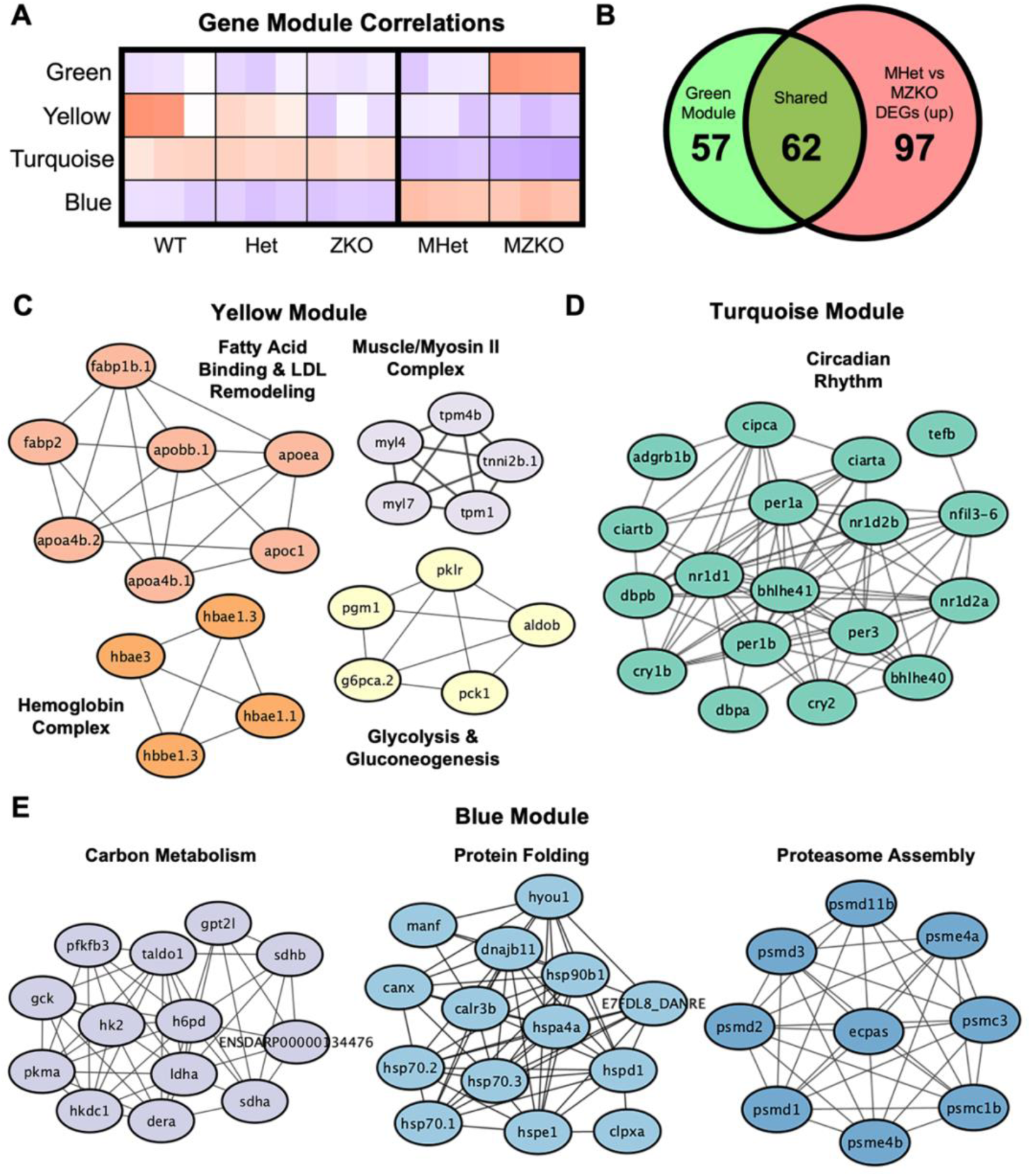
Expression signatures derived from maternal and zygotic genotypes. **A**: Heat map of modules identified through weighted gene co-expression network analysis that were highly correlated with either the maternal or zygotic genotype at 5 dpf. **B**: Overlap of genes within the green module and upregulated DEGs in MZKOs compared to their MHet siblings at 5 dpf. **C**: STRING analysis of genes within the yellow module showing interacting networks of proteins involved in fatty acid binding and LDL remodeling, glycolysis and gluconeogenesis, and the hemoglobin complex. **D**: STRING analysis of genes within the turquoise module showing an interacting network of proteins involved in circadian rhythm regulation. **E**: STRING analysis of genes within the blue module showing interacting networks of proteins involved in processes such as protein folding, carbon metabolism, and proteasome assembly.

The green module was strongly linked to disease progression, as these genes were positively correlated with the MZKO genotype, and many of these overlapped with the upregulated DEGs in MZKOs described above (**Fig. 8B**). The yellow module, in contrast, suggested shared dysregulation due to loss of *pomgnt2*, whether maternal, zygotic, or both. It included 167 genes that were negatively correlated and downregulated across ZKOs and both genotypes obtained from KO x Het crosses. This module showed enrichment of smaller STRING networks involved in glycolysis and gluconeogenesis, the hemoglobin complex, and lipid metabolism and LDL remodeling (**Fig. 8C**). The LDL network was driven by reduced *apolipoprotein* expression, similar to what was observed in MZKOs at 10 dpf indicating early dysregulation in lipid metabolism in ZKOs and their offspring. A few genes involved in muscle contraction were also present in the yellow module, including *myosin light chain 4 and 7* (*myl4, myl7*), *tropomyosin 1 and 4b* (*tpm1, tpm4b*), and *troponin I 2B* (*tnni2b.1*) (**Fig. 8C**). This module suggested overall that certain differences in physiological processes may persist from *ZKO* females to their oocytes.

The turquoise and blue modules, which were respectively negatively and positively correlated with both MZKOs and their MHet siblings, revealed a difference between offspring of Het females and ZKO females. Within the downregulated module (turquoise), numerous genes forming the structural constituents of different tissues were identified. These included multiple *crystallin* genes, which form the lens of the eye; *collagens*, the structural components of the ECM; *tubulins,* intermediate filaments, and genes involved in actin monomer binding that are critical for cytoskeletal assembly (**Supplemental Fig. 5A**). Interestingly, key regulators of circadian rhythm were also included in this module, including *per1a, per1b, per3, nr1d1, nr1d2a, nr1d2b, cry1b,* and *cry2* (**Fig. 8D**). Within the entire turquoise module, *per1b* and *nr1d1* were the most centrally connected hub genes (**Supplemental Fig. 5B**), suggesting potential drastic differences in regulation of circadian rhythm stemming from the maternal state. The upregulated module (blue), in contrast, showed increased activation of several processes including protein folding with upregulation of heat shock proteins and other chaperones, proteasome assembly, and carbon metabolism, as demonstrated by upregulation of several glycolytic and Krebs cycle enzymes in the mitochondria (**Fig.8E**). Of particular interest for its implications in maternal compensation was *hexokinase domain-containing 1* (*hkdc*), whose human ortholog has been strongly linked to glucose intolerance during pregnancy [45, 46].

Overall, our findings show how removal of maternal *pomgnt2* mRNA in oocytes is necessary to unmask developmental phenotypes that recapitulate the core dystroglycanopathy phenotypes. However, our transcriptomic analyses revealed significant differences in several physiological processes correlated with maternal genotype. Based on these data, we propose a model in which metabolic dysfunction stemming from progressive muscle disease in ZKO females results in metabolic rewiring in the offspring to balance differences in lipid and cholesterol synthesis with an increased metabolic requirement to maintain homeostasis (**Fig. 9**).

**Figure 9.**
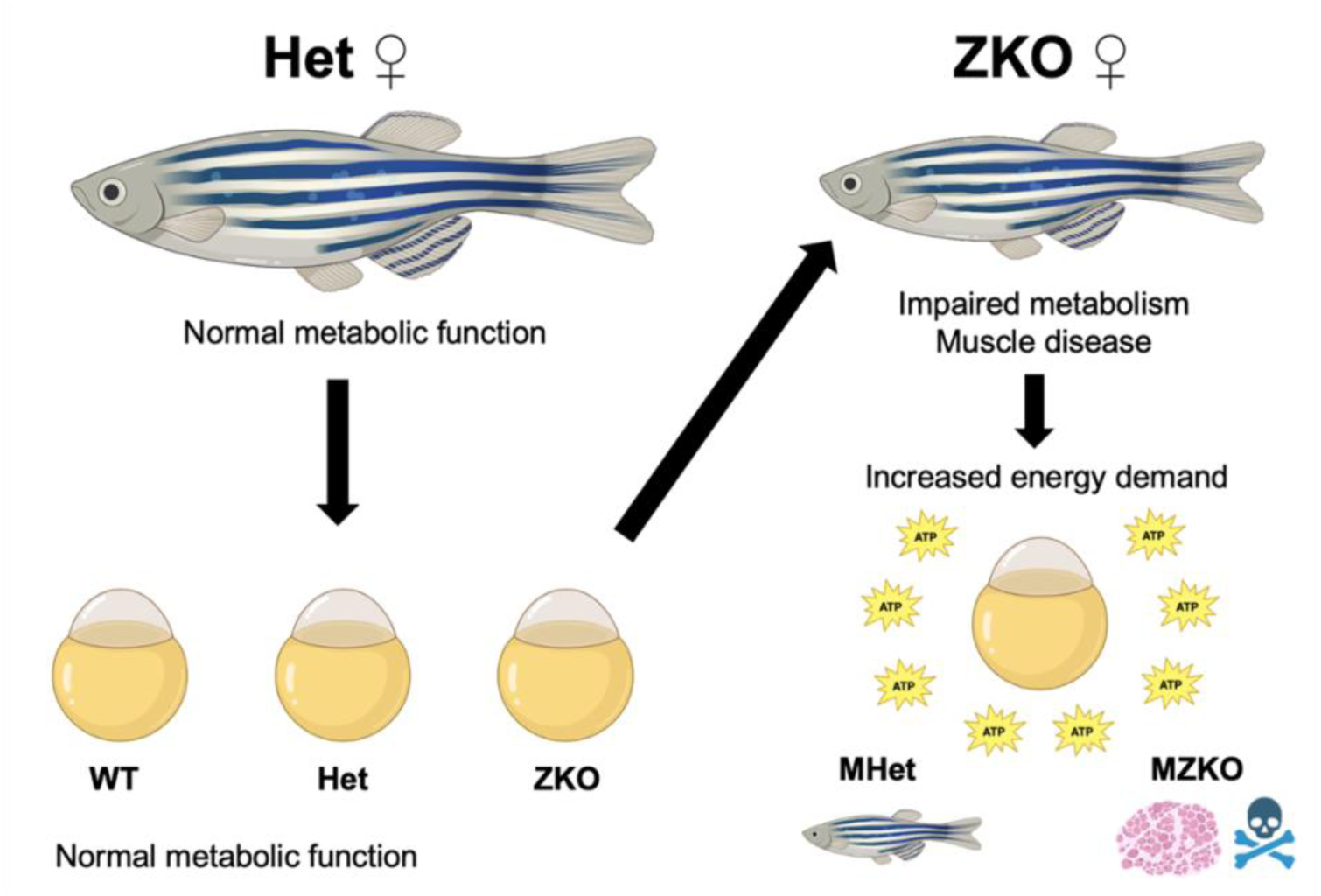
Model of metabolic differences in progeny of Heterozygous and ZKO females. Figure made with biorender.com.

## DISCUSSION

In this study, we characterized a novel zebrafish mutant for the glycosyltransferase Pomgnt2 to model severe neuromuscular disorder caused by loss α-DG glycosylation, which mediated cell-ECM interactions critical for morphogenesis and maintenance of multiple tissues [20, 22, 29, 47]. We found that *pomgnt2* is maternally provided in zebrafish, resulting in residual α-DG glycosylation in ZKO larvae masking developmental phenotypes. By generating MZKOs, we achieved complete loss of α-DG glycosylation in the embryo, and recapitulated muscle, eye, and axon guidance deficits found in severe dystroglycanopathy, also identifying molecular signatures of disease progression and pathophysiology [31, 32, 35].

Since ZKOs only show progressive muscle disease in adulthood, this line offered the additional opportunity to investigate the impact of using maternal zygotic mutants for modeling disease in zebrafish. Zebrafish offers significant advantages as a human disease model due to the conservation of many human genes and pathogenic mechanisms, especially for developmental disorders as the dystroglycanopathies [3, 48, 49]. However, genetic compensation through different mechanisms, including maternal factor deposition in the oocyte, has led to significant disparities between morphant and KO disease models [1, 4–6, 9, 50–52]. Maternal transcripts deposited in the yolk are often essential for development, making maternal-zygotic mutants necessary to unmask disease-relevant phenotypes in the zebrafish [12, 53–56]. However, not as much attention has been given to understanding the impact of the KO mother on the offspring [13, 14, 57, 58].

### The need for using maternal-zygotic mutants to study dystroglycanopathies

Zebrafish has become a valuable disease model for dystroglycanopathies due to fundamental differences in embryonic development between humans and rodents. Mouse KOs for dystroglycan (*Dag1*) and for the glycosyltransferases initiating O-mannosylation, *Pomt1* and *Pomt2*, show early embryonic lethality due a rodent-specific disruption in Reichert’s membrane during placental formation [24–26]. Thus, zebrafish KOs for *dag1*, *pomt1*, and *pomt2* are so far the only vertebrate animals to model global developmental loss of these genes.

While zebrafish *dag1* ZKOs show early lethality and progression of muscle disease consistent with morphants for other dystroglycanopathy genes [31], *pomt2* ZKOs displayed muscle, brain, and eye phenotypes only after 2 months of age [34]. These findings were reconciled by studies on *pomt1* KOs showing that later disease onset was due to maternal compensation that could be removed by generating maternal-zygotic mutants [35]. The findings in this study confirmed that maternal compensation also occurs in *pomgnt2* ZKO zebrafish, as the phenotypes identified in *pomgnt2* MZKO are not only consistent with *pomt1* MZKOs, but also with *dag1* and *fkrp* ZKOs that do not undergo maternal compensation. This will likely extend to other zebrafish models of dystroglycan function. For example, *pomgnt1* KOs have only shown retinal degeneration at 6 months post fertilization while mouse models have phenotypes consistent with dystroglycanopathy [33]. Transcripts for *pomt2*, *pomgnt1* and most other glycosyltransferases involved in α-DG glycosylation are detected in zebrafish zygote before the MZT, strongly suggesting maternal compensation will be present [37].

### Lipid metabolism disruptions due to *pomgnt2* loss of function

While *Pomgnt2* KO pups reach term, they die within the first postnatal day and only cortical migration deficits have been investigated [29]. No characterization of muscle and eye deficits has been performed, and little is known about disease progression in *POMGNT2*-related dystroglycanopathy due to the limited number of reported variants. Our developmental analysis of muscle integrity showed that *pomgnt2* MZKOs show progressive detachment of muscle fibers from the MTJs and degeneration of the slow-twitch fiber layer as muscle integrity is lost, similar to DMD (*dmd-sapje*) and merosin-deficient CMD (*lama2-candyfloss*) zebrafish models [59]. Similar phenotypes had been also noted in a morphant model for *fkrp* [60]. The rapid fiber degeneration in our model is more similar to DMD models, while detached fibers in the *lama2* zebrafish KO can survive for days following detachment [61], suggesting shared pathophysiology among mutants in the dystrophin/glycoprotein complex.

Transcriptomic analyses at 5 and 10 dpf revealed initial compensatory responses when muscle integrity is relatively preserved at 5 dpf, followed by widespread reduction in muscle contraction genes during severe disease at 10 dpf. Of particular interest were alterations in genes controlling lipid metabolism and sterol biosynthesis. Dyslipidemia and other disruptions in lipid metabolism are characterized in DMD [62–64]. Studies on individuals with DMD and animal models not only show different profiles of cholesterol, phospholipids, and fatty acids in dystrophic muscle and serum, but also identified changes in membrane phospholipid composition and fatty acid metabolism in mitochondria that actively contribute to muscle disease progression [65–67]. Much less is known about these processes in dystroglycanopathies. One metabolomic study using a knock-in *Fkrp* line (*Fkrp^P448L^*) to model a less severe form of dystroglycanopathy, Limb Girdle Muscular Dystrophy 2i (LGMD2i), identified global metabolic perturbations with increases in glycolytic intermediates and lipid metabolites [68]. Our findings in 10 dpf *MZKO*s demonstrate broad downregulation of genes involved in cholesterol, phosphoglyceride, and lipoprotein metabolism, again suggesting shared pathophysiology with DMD that warrants further investigation.

### Implication of maternal KO status

To our knowledge, this study provides the first direct transcriptomic comparison of larvae from Het X Het and KO X Het crosses. WGCNA analysis revealed clear differences associated with maternal genotype, highlighting important consideration specific to generating maternal-zygotic KOs through zygotic KO females. Offspring of ZKO females showed decreased expression of structural genes (collagens, tubulins, and crystallins), suggesting potential differences in tissue formation and development. In addition, the reduction in circadian regulators warrant further investigation into possible differences in rhythmicity.

On the other hand, increased expression of genes involved in protein homeostasis and glycolysis suggest that offspring of ZKO mothers may be primed for an increased metabolic requirement to maintain normal cellular function. Despite sharing these gene expression differences with their MZKO siblings, MHet are behaviorally and histologically undistinguishable from WTs and Hets from HetxHet crosses. Thus, while this increased expression of metabolic genes may not have major deleterious impact on the overall health of the offspring, it could modify disease progression in MZKOs. Some metabolic gene expression changes were noted in ZKOs as early as 5 dpf when they still benefit from residual α-DG glycosylation. It is possible that ZKOs have life-long changes that compound with progressive muscle disease in adult females with consequences for the maternal nutrition and metabolism altering the yolk of the oocytes, which is rich in lipids and proteins [69, 70]. Several studies have also noted reduced offspring viability and other abnormalities in zebrafish in response to metabolic disruptions in the female parents [62, 71, 72]. These differences could be leveraged in future studies to identify novel disease modifiers.

In conclusion, our study shows how a maternal-zygotic mutant can be leveraged to unveil early disease phenotypes in strains impacted by maternal compensation, while highlighting physiological differences that emerge when offspring are obtained from zygotic mutant females. These findings have immediately relevance for other zebrafish mutants of dystroglycan-related disorders as well as other zebrafish mutants where maternal compensation is suspected.

## MATERIALS AND METHODS

### Experimental Model

All experiments and procedures involving live animals in this study were approved by the Institutional Animal Care and Use committee of Rutgers University. Zebrafish were housed on a recirculating Tecniplast USA system under a 14-10 light-dark cycle at 28°C in 3.5 L tanks and fed twice daily. For spawning events, males and females were place off-system in divided spawning cages each evening, and the dividers were removed the following morning following the start of the lights on period. Embryos obtained from spawning events were collected in petri dishes with egg water containing methylene blue (5×10^-5^ % w/v) until 5-6 days post fertilization (dpf) when they were placed on the system for raising or until their respective endpoints.

### Guide RNA (gRNA) assembly

Three guide RNA (gRNA) target sequences were identified from the Burgess Lab UCSC Track Data Hub for CRISPR targets and ordered from Integrated DNA Technologies (IDT; Coralville, IA) with SP6 or T7 promoter sequence, gRNA target sequence, and TracrRNA overlap sequencing. 10 µM of each gRNA oligo and 10 µM of universal oligo were annealed in 25 µl reactions with 1U Phusion™ High Fidelity DNA Polymerase (Thermo Fisher Scientific) in a T100™ Thermal Cycler (Bio-Rad) under the following cycling conditions: 98°C for 2 minutes, 50°C for 10 minutes, and 72°C for 10 minutes. gRNAs were then synthesized using a HiScribe SP6 or T7 Quick High Yield RNA Synthesis kit (New England Biolabs) in a 30 µl containing 3 µl of annealed oligo product at 37°C overnight. gRNAs were then purified using an RNA Clean & Concentrator kit (Zymo Research), diluted in sterile, nuclease free water, and stored at - 80°C for no longer than one month.

### Generation of the *pomgnt2* line

Microinjections were performed on one cell stage EK zebrafish embryos using 1 nanoliter/embryo of injection master mix containing approximately 90 pg/nl of each gRNA and 500 pg/nl recombinant Cas9 protein (PNA Bio Inc). Injections using a gRNA targeting *tyrosinase* (*tyr*) was used to determine technique efficiency for each experiment by screening the developing embryos for loss of pigment. Uninjected clutchmate controls were also used in every experiment. The injected embryos were housed in embryo medium with methylene blue and screened periodically for 5 days to remove dead and deformed embryos, and the remaining were placed on the system to be raised for mutant line propagation. Between 3-4 months post fertilization (mpf), surviving injected zebrafish were anesthetized in 0.016% w/v tricaine methane sulfonate (Tricaine, MS-222) and fin clipped. DNA was extracted and heteroduplex mobility assays were used to identify fish harboring indels successfully induced through CRISPR-Cas9. The fish demonstrating the highest degree of heteroduplex formation were then outcrossed with WT EK zebrafish to generate F_1_ founders.

### F_1_ founder selection

When potential F_1_ founders reached 3-4 months of age, they were anesthetized and fin clipped, and DNA was extracted and amplified from fin clips as previously done for the F_0_ generation. PCR products and forward primers for each reaction were sent to Azenta Life Sciences (South Plainfield, NJ) for Sanger Sequencing. Five F_1_ founders (4 female, 1 male) with a 13 bp insertion and 4 bp deletion in exon 1 along with a 7 bp deletion in exon 2 were selected to propagate the main mutant line described in this study. Additional founders with only the exon 1 mutations or only the exon 2 mutations were identified and later used to corroborate loss of pomgnt2 function. F_1_ founders were then outcrossed again to WT EK fish to further propagate the line.

### Genotyping of the pomgnt2 line

Following the identification of the primary set of F_1_ founder mutations, we used several validated methods to genotype the fish for experiments. For the HetxHet crosses, DNA was extracted and amplified exactly as done for the F_0_ and F_1_ generation. Following PCR amplification, 10 µl of PCR product was digested at 37°C for 1-2 hours to overnight with 1U SnaBI restriction enzyme, which recognizes the sequence that is deleted in exon 2 (New England Biolabs) in a 25 µl reaction. Digestion was then stopped by incubating the samples at 4°C. Each sample was run on a 1.5% agarose gel at 90 V for 1 hour and imaged under UV light to visualize banding pattern. Random samples were also periodically spot checked via Sanger Sequencing using amplicons of both the exon 1 and 2 mutations to further ensure genotyping accuracy. Genotyping was also performed through quantitative allele-specific PCR with custom designed Affinity Plus™ qPCR probes and primers from Integrated DNA Technologies (Coralville, IA) on a QuantStudio™ 6 qPCR system (Applied Biosystems) for 40 cycles with an annealing temperature of 62°C. Due to differences in WT and mutant probe efficiency, qPCR-based genotyping reactions were often run with a second reaction containing only the WT probe, along with multiple controls genotyped through alternate methods (i.e. restriction digest or Sanger sequencing) to ensure genotypic accuracy.

### Survival and morphological analysis

Progeny of Heterozygous (HetxHet) crosses were placed on system at 5 dpf ungenotyped with reduced water flow as close to maximum allowable stocking densities as possible (5-30 dpf: 14/L; >30 dpf: 4/L) to induce competition for food. The same process was performed for KOxHet survival and morphological analyses with a slightly reduced initial stocking density to accommodate smaller clutches generated from KO females (11-13/L). At regularly schedule intervals of 1 month, 3 months, and 5 months for HetxHet crosses and 7, 10, 14, and 28 days for KOxHet crosses, the animals were removed from the system, imaged to obtain body length measurements, and either fin clipped and returned to the system for breeding and other experimental purposes, or sacrificed for other experimental purposes. At timepoints of 1 month of less, the juveniles were imaged in 3% methylcellulose with a M165 FC stereo microscope and LAS Software v4.21 (Leica Microsystems). At 3 and 5 months, the fish were imaged with a handheld camera to obtain body length measurements. Body length measurements were taken from the mouth to the base of the tail fin in ImageJ with the researcher masked to genotype.

### RNA isolation

Total RNA was extracted with RNAzol® RT RNA Isolation reagent as follows: Each sample was homogenized with 500 µl of RNAzol. 200 µl of nuclease free water was added to each sample, which was then mixed vigorously for 15 seconds and incubated at room temperature for 15 minutes. The samples were then centrifuged at 12,000 x g for 15 minutes at 4°C. 600 µl of supernatant was then removed and added to a fresh 1.5 ml Eppendorf tube with 600 µl of 100% isopropanol and mixed by inverting. Samples were then incubated at −80°C for 30 minutes, then at room temperature for 15 minutes, and then centrifuged at 12,000 x g for 10 minutes at 4°C. The supernatant was then removed, and the pellet was washed with 1 ml of 75% ethanol and centrifuged at 8,000 x g for 3 minutes at 4°C. The supernatant was removed, and the pellet was washed in 100 µl of 75% ethanol 3-4 more times with centrifuging in between. The pellet was then dried on a heat block at 50°C, resuspended in nuclease free water, and stored at −80°C for used in downstream applications.

### Quantitative reverse transcriptase PCR (qRT-PCR)

qRT-PCR was performed to evaluate *pomgnt2* gene expression across a time course in HetxHet crosses at 20, 40, 60, and 90 dpf in composite samples of genotyped heads (20, 40, and 60 dpf) or tails (90 dpf). Following RNA isolation, 1 µg of RNA was reverse transcribed using iScript™ Reverse Transcription Supermix (Bio-Rad) in a 20 µl reaction containing 16 µl of RNA and 4 µl of RT. cDNA was then diluted from 50 ng/µl to 20 ng/µl in nuclease free water. qPCR reactions were run using PowerUp™ SYBR™ Green Master Mix (Applied Biosystems) in triplicate 15 µl reactions with final cDNA and primer concentrations of 10 ng/ul and 400 nM, respectively, on a QuantStudio™ 3 qPCR system (Applied Biosystems) for 40 cycles with annealing temperatures of 56°C for the *pomgnt2* and 53°C for *rpl13α*, the endogenous control. Each experiment was run with no template controls and 1-2 no RT controls per primer pair. Each datapoint is presented as fold change (2^-ΔΔCT^) compared to the average of the WT samples.

### Western blotting and WGA enrichment

The western blotting procedure used in this study was previously described by Karas et al. [35]. Western blotting was performed on composite samples of 7-10 30 dpf fish using wheat germ agglutinin (WGA) enrichment of glycoproteins, as previously described by Karas et al. Composite samples were lysed in 100 µl of buffer made in-house (50 mM Tris pH 8, 100 mM NaCl, 1 mM PMSF, 1 mM Na Orthovanadate, 1% Triton-X 100), centrifuged at 4°C for 30 minutes, sonicated 3 times for 5 minutes, and treated with 2 µl DNase I (New England Biolabs). Protein concentration in each lysate was quantified using a BCA Protein Assay kit (G-Biosciences). 500 µg of protein was diluted in 200 µl of lectin binding buffer (20 mM Tris pH 8, 1 mM MnCl_2_ and 1 mM CaCl_2_), added to 50 µl of WGA agarose-bound beads (Vector Laboratories), and incubated overnight, rocking at 4°C. The following morning, the samples were centrifuged at 15,000 x g for 2 minutes to remove the supernatant, and the bead-bound glycoproteins were eluted with 30 µl of 4X Laemmli buffer (Bio-Rad). Glycoproteins were separated on a 4-12% Bis-Tris Protein SDS-PAGE gel (Life Technologies) run at 55 V for 10 minutes, then 120 V for 70 minutes, in 1X Invitrogen™ NuPAGE™ MOPS SDS Running Buffer. The gel was then incubated in transfer buffer (70% Water, 20% Methanol, 10% 1X Tris/Glycine Buffer with 0.1% SDS (Bio-Rad) for 5-10 minutes. Proteins were then transferred to a nitrocellulose membrane (Thermo Fisher) at 4°C in transfer buffer with an ice pack at 110 V for 3.5 hours. The membranes were then stained with 0.1% naphthol blue black (amido black) (Millipore Sigma) for total protein staining. Nitrocellulose membranes were blocked in 5% milk diluted in 1X TBS containing 0.1% Tween-20 (TBS-T), then probed with 1:100 anti-α-dystroglycan antibody clone IIH6C4 (Millipore Sigma) and 1:1000 anti-*β*-dystroglycan antibody (ab62373, Abcam) overnight at 4°C. The following day, the membranes were briefly rinsed in MilliQ water, probed with 1:3000 peroxidase AffiniPure donkey anti-mouse IgG and 1:20 000 peroxidase AffiniPure donkey anti-rabbit IgG (Jackson ImmunoResearch), washed in 1X TBS-0.1% Tween-20 5X for 5 minutes, and developed with chemiluminescence Pierce ECL Western Blotting Substrate on CL-XPosure™ Film (Thermo Fisher).

### Automated behavior tracking in larvae

Automated tracking of locomotor behavior in larvae was performed using a DanioVision™ Observation Chamber (Noldus Information Technology, Wageningen) and EthoVision® XT Video Tracking software at 30 frames per second. All experiments were done in a randomized, ungenotyped manner. At 5 dpf, larvae were placed in a 96 well polystyrene cell culture plate in equal water volumes and placed in the observation chamber for a total of one hour: the first 30 minutes for habituation, and the last 30 minutes for tracking. At 7, 10, and 14 dpf, larvae were placed in a 24 well polystyrene cell culture plate to increase the total area for movement. These experiments were performed in the same manner, but with 20 minutes of habituation and 20 minutes of tracking. The observation chamber was always held at a constant 28.5°C and experiments always began at the same time each day to minimize variability.

### Behavior tracking in adult fish

Adult WT and ZKO fish were evaluated for locomotor function at approximately one year of age. Each fish had been genotyped prior to the experiment, which occurred over the course of four days, evaluating one fish at a time in alternating WT-ZKO order whenever possible. The experiments always began at the same time each day to minimize variability between experiment days. Each fish was placed in an arena of 60 cm in diameter held at approximately 26°C ± 1.5. The fish were habituated for 5 minutes, followed by 10 minutes of recording using a GS3-U3-41C6NIR-C 1” FLIR Grasshopper video camera at 30 frames per second. The data was then analyzed using AnimalTA Tracking Software [73].

### Cryosectioning and fluorescent immunohistochemistry

All fish, larvae, juvenile, or adult, were fixed in 4% paraformaldehyde (PFA) overnight at 4°C. The fish were in 1X PBS 3 times for 5 minutes to remove any residual PFA. The fish were then transferred to 15% sucrose with 0.2% sodium azide in 1X PBS for 1-2 days, followed by 30% sucrose with 0.2% sodium azide in 1X PBS for 1-2 days, and then frozen in isopentane on dry ice in Tissue Freezing Medium (Ted Pella Inc) and stored at −80°C. Fixed tissue from adult fish was also decalcified for 1-2 hours in Cal-Ex (Fisher Scientific) and washed in 1X PBS 3 times for 5 minutes before beginning cryoprotection in sucrose. Cryosectioning was performed on a Leica CM1850 UV Cryostat (Leica Microsystems) generating 12 µm transverse sections of the eyes and brain in larvae and 16 µm transverse sections of muscle in adults. The sections were dried on a slide warmer and either stained immediately or stored at −20°C until staining.

For transverse sections of adult muscle, sections were outlined, blocked in 10% normal goat serum (NGS) containing 1% Triton X 100 and 2% Tween-20 in a humidified slide box at room temperature for 1 hour. Next, the sections were incubated in primary antibody diluted in 1% NGS containing 1% Triton X 100 and 2% Tween-20 overnight at 4°C. The primary antibodies used in these experiments were 1:50 anti-laminin L9393 (Millipore Sigma) and 1:50 anti-DAG1 antibody clone IIH6C4 (Abcam). The following day, the sections were washed in 1X PBS 3 times for 5 minutes and incubated in secondary antibody diluted in 1% NGS containing 1% Triton X 100 and 2% Tween-20 at room temperature for one hour. Alexa Fluor™ goat anti-rabbit and goat anti-mouse secondaries (Thermo Fisher) were used in these experiments at 1:250 concentration. Sections were then counterstained in 1X DAPI or Hoechst for 10 minutes, coverslipped with VWR micro cover glasses in Prolong™ Gold Antifade Mountant (Thermo Fisher) and cured for at least 24 hours before imaging.

For transverse sections of the eyes and brain in larvae, this process was repeated with the following modifications: blocking was performed in 10% NGS with 1% Triton X 100, primary and secondary antibodies were diluted in 1% NGS with 0.1% Triton X 100, and counterstaining in 1X DAPI was done for 1 hour. The primary antibodies used in these experiments were 1:200 anti-synaptophysin (Abcam), 1:200 zpr1 (ZIRC), and 1:100 zn-8 (Developmental Studies Hybridoma Bank). The secondary antibodies and dilutions were unchanged. All cryosections were imaged on a Zeiss LSM800 confocal microscope with Zeiss Zen imaging software.

### Whole mount fluorescent immunohistochemistry

The protocol for whole mount staining was adapted from Bailey et al. [74]. Fluorescent immunohistochemistry to analyze muscle in larvae was performed through a whole mount staining protocol adapted from Bailey et al. Zebrafish fixed overnight in 4% PFA at 4°C. The following day, they were rinsed 3 times for 10 minutes in 1X PBS-0.1% Tween 20 (PBS-T), followed by permeabilization in 1 mg/ml collagenase D (Sigma) for 1.5 hours at room temperature. The fish were then washed 3 times for 10 minutes in PBS-T. For experiments using Alexa Fluor™ Phalloidin 546 (Thermo Fisher) and/or Alexa Fluor^™^ 488 α-bungarotoxin conjugate (Invitrogen), these steps were performed at this stage at 1:20 and 1:500 dilutions, respectively, for 2 hours at room temperature, followed by additional washes in PBS-T. The fish were then blocked overnight in antibody blocking solution (Ab block) made in-house (5% BSA, 1% DMSO, 1% Triton-X-100, 0.2% saponin in 1X PBS) at 4°C. The following day, the fish were moved to primary antibody diluted in Ab block. The primary antibodies used in these experiments were 1:50 anti-DAG1 antibody clone IIH6C4 (ab234587, Abcam), 1:50 anti-laminin L9393 (Millipore Sigma), 1:25 F59 (ZIRC), and 1:10 anti-SV2 (Developmental Studies Hybridoma Bank). Primary antibody incubations were performed overnight at 4°C. The following day, the fish were then washed out of primary in PBS-T 3 times for 10 minutes each and moved to secondary antibody diluted in Ab block. 1:250 Alexa Fluor^™^ goat anti-mouse and goat anti-rabbit secondaries (Thermo Fisher) were used for these experiments. The following day, the fish were washed out of secondary in PBS-T, mounted in 1% low melt agarose, and imaged on a Zeiss LSM800 confocal microscope with Zeiss Zen imaging software.

### RNA-sequencing and analysis

RNA sequencing was performed by Novogene Corporation (Sacramento, CA) on RNA extracted from composite samples of 4-6 whole larvae at 5 and 10 dpf. At 4 dpf, DNA was extracted from live progeny of HetxHet and KOxHet crosses using a Zebrafish Embryonic Genotyper (ZEG) Microfluidic system (wFluidx Inc) and genotyped directly using our custom qPCR assay. At 5 dpf, the fish were sorted by genotype, euthanized in Tricaine, and snap frozen in liquid nitrogen. Fish used for the 10 dpf experiment were housed on system separated by genotype until this timepoint. Total RNA was extracted using RNAzol® RT RNA Isolation reagent as described in the **RNA isolation** section and shipped overnight on dry ice for sequencing. In addition, leftover RNA from the composite samples was reverse transcribed using iScript™ Reverse Transcription Supermix (Bio-Rad) as described in the “Quantitative reverse transcriptase PCR (qRT-PCR)” section and genotyped again using our Affinity Plus^TM^ qPCR probes to ensure all fish were sorted correctly.

143.6 Gb of raw data were delivered as fastq files with an average of 44.5 million raw reads for the 5 dpf experiment and 48.4 million reads for the 10 dpf experiment, which were uploaded to Amarel, the high-performance computing cluster of Rutgers University. Quality control was performed in house using FastQC [75]. Following FASTQC warnings in Per Base Sequence Content, the first 10 bp of each read was trimmed to remove residual adapter sequence bias and reads with Phred scores <20 were filtered out before alignment. Filtered sequencing reads were aligned to the GRCz11 (danRer11) genome build with HISAT2 [76] followed by sorting and indexing through Samtools [77], each of which were performed using default settings. Filtered sequencing reads were aligned to the GRCz11 (danRer11) genome build with HISAT2 [76] followed by sorting and indexing through Samtools [77]. Alignment statistics for each sample are detailed in **Supplemental Table 2**. Mapped reads were counted from sorted bam files using the featurecounts command of the Subread package in R [78] using the Lawson Lab Zebrafish Transcriptome Annotation v4.3.2 [79] as an index to improve mapping through well-defined 3’UTR annotations. Count matrices were then normalized through DESeq2 [80] and differential expression was calculated across genotypes with a adjusted p-value <0.01 and fold change >0.7 or <-0.7 to determine significance. Enrichment analysis was performed in STRING through manual curation using a combination of available databases including Gene Ontology, Zebrafish Phenotype Ontology, STRING Local Networks, COMPARTMENTS, and Pfam and Interpro Protein Domains. All sequencing data is available from the corresponding author upon request.

### Weighted gene co-expression network analysis

Following differential expression analysis, weighted gene co-expression network analysis (WGCNA) [81, 82] was performed in RStudio on 5 dpf RNA sequencing reads in WT, Het, ZKO, MHet, and MZKO samples. This timepoint was selected to perform a direct comparison between HetxHet and KOxHet progeny at a timepoint when residual α-dystroglycan glycosylation was still present in *MZpomgnt2s*. The data was variance stabilized and reduced to the 95^th^ quantile leaving a total of 1818 genes remaining. Clustering was performed using Scale Free Topology Model Fit with a soft thresholding power of 12. The genes were then clustered into modules of correlated genes which were prioritized based on correlations with both the female parent’s genotype and the progeny’s genotype. To construct the networks for each module, edge lists were generated from topological overlap matrices (TOMs) using a minimum correlation threshold of 0.4. The top 50 edges were filtered and visualized in Cytoscape [83] to identify the most centrally connected hub genes for each module.

### Quantitative analysis of neuromuscular junctions

Maximum intensity projections of the NMJs, marked by SV2 and α-bungarotoxin (α-BTX), were generated from z-stacks taken using identical imaging parameters at 20X magnification with a scan area of 638.9 µm x 638.9 µm. For each slice, the sample area was scanned four times and averaged with 1 µm intervals. The images were processed in Fiji and imported into a custom CellProfiler [84] pipeline. 3-4 hemi-segments and myotendinous junctions (MTJs) were traced per fish. The SV2 channel was rescaled identically across all samples when analyzing muscle fibers within the hemi-segments to minimize background fluorescence, but this was not necessary when analyzing MTJs. α-BTX and SV2 puncta were detected using Otsu’s thresholding method with three classes. Pixel intensity was recorded for each puncta and averaged across each fish, and colocalization was quantified using Pearson’s correlation coefficient. All NMJ images were acquired and analyzed with the researcher masked to genotype.

### Quantitative analysis of photoreceptor synapses

Raw czi files from cryosections of the entire retina were taken at 20X magnification with identical imaging parameters. The outer plexiform layer was traced in CellProfiler from the dorsal ciliary margin to the optic nerve exit without rescaling or any other modification to the image, and average pixel intensity was obtained. The images were analyzed with the researcher masked to genotype.

### Statistical analysis

All statistical analyses, aside from RNA sequencing analyses, was performed in GraphPad Prism v.8.20 (GraphPad, San Diego, CA). Normality of each dataset was assessed using a Shapiro-Wilk test. For comparisons between 2 groups of a single measure, an unpaired t-test was used, or alternatively a Mann-Whitney test for nonparametric datasets. A two-way ANOVA was used for comparisons between 2 or more groups across multiple coniditons (i.e. timepoint). Statistical significance was defined as a p-value <0.05 (*<0.05; **<0.01; ***<0.001; ****<0.0001). For all experiments except for survival analyses, each datapoint represents an individual fish (n). For survival analyses, each datapoint represents the percent genotype of an independent clutch of fish (N), while total number of individual fish (n) per clutch is listed in each figure legend.

## ACKNOWLEDGEMENTS

We are incredibly grateful to Kathleen Flaherty of the Rutgers Zebrafish Facility and Rutgers Animal Care staff members for their dedicated work in zebrafish care and the Office of Advanced Research Computing at Rutgers for high performance computing access and maintenance. We also thank our many colleagues and collaborators for insightful discussion on experimental approaches and analyses, including Dr. Clarissa Henry (University of Maine) and Dr. Ronald Hart (Rutgers University).

## AUTHOR CONTRIBUTIONS

K.P.F., N.B., B.F.K., and K.R.T. generated and propagated the *pomgnt2* strain. K.P.F., S.M., N.B., L.R.C., C.D.O., C.V., and L.M.S. performed the experiments. K.P.F., D.L., D.B.L., and M.C.M. analyzed the data. K.P.F. and M.C.M. wrote the manuscript. All authors approved of the final manuscript.

## DATA AVAILABILITY STATEMENT

All data generated in this study is available from the corresponding author upon request.

## CONFLICT OF INTEREST STATEMENT

The authors declare no conflicts of interest.

## FUNDING

This work was funded by the National Institute of Neurological Disorders and Stroke (R01NS109149) to M.C.M., the Robert Wood Johnson Foundation (grant #74260) to M.C.M., and a CureCMD Pilot Grant to M.C.M.

## Notes

### Competing Interest Statement

The authors have declared no competing interest.

## REFERENCES

1. Rouf, M.A., et al., The recent advances and future perspectives of genetic compensation studies in the zebrafish model. Genes Dis, 2023. 10(2): p. 468–479.

2. Glasauer, S.M. and S.C. Neuhauss, Whole-genome duplication in teleost fishes and its evolutionary consequences. Mol Genet Genomics, 2014. 289(6): p. 1045–60.

3. Howe, K., et al., The zebrafish reference genome sequence and its relationship to the human genome. Nature, 2013. 496(7446): p. 498–503.

4. Cardenas-Rodriguez, M., et al., Genetic compensation for cilia defects in cep290 mutants by upregulation of cilia-associated small GTPases. J Cell Sci, 2021. 134(14).

5. Nord, H., et al., Genetic compensation between Pax3 and Pax7 in zebrafish appendicular muscle formation. Dev Dyn, 2022. 251(9): p. 1423–1438.

6. Kok, F.O., et al., Reverse genetic screening reveals poor correlation between morpholino-induced and mutant phenotypes in zebrafish. Dev Cell, 2015. 32(1): p. 97–108.

7. Karakas, B., et al., P21 gene knock down does not identify genetic effectors seen with gene knock out. Cancer Biol Ther, 2007. 6(7): p. 1025–30.

8. Moreno, R.L., et al., Investigation of Islet2a function in zebrafish embryos: Mutants and morphants differ in morphologic phenotypes and gene expression. PLoS One, 2018. 13(6): p. e0199233.

9. Rossi, A., et al., Genetic compensation induced by deleterious mutations but not gene knockdowns. Nature, 2015. 524(7564): p. 230–3.

10. Fraher, D., et al., Zebrafish Embryonic Lipidomic Analysis Reveals that the Yolk Cell Is Metabolically Active in Processing Lipid. Cell Rep, 2016. 14(6): p. 1317–1329.

11. Aanes, H., et al., Zebrafish mRNA sequencing deciphers novelties in transcriptome dynamics during maternal to zygotic transition. Genome Res, 2011. 21(8): p. 1328–38.

12. Lee, M.T., A.R. Bonneau, and A.J. Giraldez, Zygotic genome activation during the maternal-to-zygotic transition. Annu Rev Cell Dev Biol, 2014. 30: p. 581–613.

13. Sun, Y., et al., Craniofacial and cardiac defects in chd7 zebrafish mutants mimic CHARGE syndrome. Front Cell Dev Biol, 2022. 10: p. 1030587.

14. Hayes, M., et al., ptk7 mutant zebrafish models of congenital and idiopathic scoliosis implicate dysregulated Wnt signalling in disease. Nat Commun, 2014. 5: p. 4777.

15. Muntoni, F., et al., Defective glycosylation in muscular dystrophy. Lancet, 2002. 360(9343): p. 1419–21.

16. Muntoni, F., S. Torelli, and M. Brockington, Muscular dystrophies due to glycosylation defects. Neurotherapeutics, 2008. 5(4): p. 627–32.

17. Kanagawa, M., Dystroglycanopathy: From Elucidation of Molecular and Pathological Mechanisms to Development of Treatment Methods. Int J Mol Sci, 2021. 22(23).

18. Yoshida-Moriguchi, T. and K.P. Campbell, Matriglycan: a novel polysaccharide that links dystroglycan to the basement membrane. Glycobiology, 2015. 25(7): p. 702–13.

19. Halmo, S.M., et al., Protein O-Linked Mannose β-1,4-N-Acetylglucosaminyl-transferase 2 (POMGNT2) Is a Gatekeeper Enzyme for Functional Glycosylation of α-Dystroglycan. J Biol Chem, 2017. 292(6): p. 2101–2109.

20. Ogawa, M., et al., GTDC2 modifies O-mannosylated α-dystroglycan in the endoplasmic reticulum to generate N-acetyl glucosamine epitopes reactive with CTD110.6 antibody. Biochem Biophys Res Commun, 2013. 440(1): p. 88–93.

21. Manzini, M.C., et al., Ethnically diverse causes of Walker-Warburg syndrome (WWS): FCMD mutations are a more common cause of WWS outside of the Middle East. Hum Mutat, 2008. 29(11): p. E231–41.

22. Manzini, M.C., et al., Exome sequencing and functional validation in zebrafish identify GTDC2 mutations as a cause of Walker-Warburg syndrome. Am J Hum Genet, 2012. 91(3): p. 541–7.

23. Beltrán-Valero de Bernabé, D., et al., Mutations in the O-mannosyltransferase gene POMT1 give rise to the severe neuronal migration disorder Walker-Warburg syndrome. Am J Hum Genet, 2002. 71(5): p. 1033–43.

24. Williamson, R.A., et al., Dystroglycan is essential for early embryonic development: disruption of Reichert’s membrane in Dag1-null mice. Hum Mol Genet, 1997. 6(6): p. 831–41.

25. Willer, T., et al., Targeted disruption of the Walker-Warburg syndrome gene Pomt1 in mouse results in embryonic lethality. Proc Natl Acad Sci U S A, 2004. 101(39): p. 14126–31.

26. Hu, H., et al., Conditional knockout of protein O-mannosyltransferase 2 reveals tissue-specific roles of O-mannosyl glycosylation in brain development. J Comp Neurol, 2011. 519(7): p. 1320–37.

27. Chan, Y.M., et al., Fukutin-related protein is essential for mouse muscle, brain and eye development and mutation recapitulates the wide clinical spectrums of dystroglycanopathies. Hum Mol Genet, 2010. 19(20): p. 3995–4006.

28. Kurahashi, H., et al., Basement membrane fragility underlies embryonic lethality in fukutin-null mice. Neurobiol Dis, 2005. 19(1-2): p. 208–17.

29. Yagi, H., et al., AGO61-dependent GlcNAc modification primes the formation of functional glycans on α-dystroglycan. Sci Rep, 2013. 3: p. 3288.

30. Nakagawa, N., et al., Ectopic clustering of Cajal-Retzius and subplate cells is an initial pathological feature in Pomgnt2-knockout mice, a model of dystroglycanopathy. Sci Rep, 2015. 5: p. 11163.

31. Gupta, V., et al., The zebrafish dag1 mutant: a novel genetic model for dystroglycanopathies. Hum Mol Genet, 2011. 20(9): p. 1712–25.

32. Serafini, P.R., et al., A limb-girdle muscular dystrophy 2I model of muscular dystrophy identifies corrective drug compounds for dystroglycanopathies. JCI Insight, 2018. 3(18).

33. Liu, Y., et al., Eyes shut homolog (EYS) interacts with matriglycan of O-mannosyl glycans whose deficiency results in EYS mislocalization and degeneration of photoreceptors. Sci Rep, 2020. 10(1): p. 7795.

34. Liu, Y., et al., Deletion of POMT2 in Zebrafish Causes Degeneration of Photoreceptors. Int J Mol Sci, 2022. 23(23).

35. Karas, B.F., et al., Removal of pomt1 in zebrafish leads to loss of α-dystroglycan glycosylation and dystroglycanopathy phenotypes. Hum Mol Genet, 2024.

36. Osborn, D.P.S., et al., Mutations in INPP5K Cause a Form of Congenital Muscular Dystrophy Overlapping Marinesco-Sjögren Syndrome and Dystroglycanopathy. Am J Hum Genet, 2017. 100(3): p. 537–545.

37. White, R.J., et al., A high-resolution mRNA expression time course of embryonic development in zebrafish. Elife, 2017. 6.

38. Fallon, J.R. and Z.W. Hall, Building synapses: agrin and dystroglycan stick together. Trends Neurosci, 1994. 17(11): p. 469–73.

39. Gee, S.H., et al., Dystroglycan-alpha, a dystrophin-associated glycoprotein, is a functional agrin receptor. Cell, 1994. 77(5): p. 675–86.

40. Hopf, C. and W. Hoch, Agrin binding to alpha-dystroglycan. Domains of agrin necessary to induce acetylcholine receptor clustering are overlapping but not identical to the alpha-dystroglycan-binding region. J Biol Chem, 1996. 271(9): p. 5231–6.

41. Wright, K.M., et al., Dystroglycan organizes axon guidance cue localization and axonal pathfinding. Neuron, 2012. 76(5): p. 931–44.

42. Lindenmaier, L.B., et al., Dystroglycan is a scaffold for extracellular axon guidance decisions. Elife, 2019. 8.

43. Davison, C. and F.R. Zolessi, Slit2 is necessary for optic axon organization in the zebrafish ventral midline. Cells Dev, 2021. 166: p. 203677.

44. Morise, J., H. Takematsu, and S. Oka, The role of human natural killer-1 (HNK-1) carbohydrate in neuronal plasticity and disease. Biochim Biophys Acta Gen Subj, 2017. 1861(10): p. 2455–2461.

45. Hayes, M.G., et al., Identification of HKDC1 and BACE2 as genes influencing glycemic traits during pregnancy through genome-wide association studies. Diabetes, 2013. 62(9): p. 3282–91.

46. Guo, C., et al., Coordinated regulatory variation associated with gestational hyperglycaemia regulates expression of the novel hexokinase HKDC1. Nat Commun, 2015. 6: p. 6069.

47. Ogawa, M., et al., N-acetylglucosamine modification in the lumen of the endoplasmic reticulum. Biochim Biophys Acta, 2015. 1850(6): p. 1319–24.

48. Veldman, M.B. and S. Lin, Zebrafish as a developmental model organism for pediatric research. Pediatr Res, 2008. 64(5): p. 470–6.

49. Moore, C.J., H.T. Goh, and J.E. Hewitt, Genes required for functional glycosylation of dystroglycan are conserved in zebrafish. Genomics, 2008. 92(3): p. 159–67.

50. Stainier, D.Y.R., et al., Guidelines for morpholino use in zebrafish. PLoS Genet, 2017. 13(10): p. e1007000.

51. She, J., et al., Genetic compensation by epob in pronephros development in epoa mutant zebrafish. Cell Cycle, 2019. 18(20): p. 2683–2696.

52. Buglo, E., et al., Genetic compensation in a stable slc25a46 mutant zebrafish: A case for using F0 CRISPR mutagenesis to study phenotypes caused by inherited disease. PLoS One, 2020. 15(3): p. e0230566.

53. Dosch, R., et al., Maternal control of vertebrate development before the midblastula transition: mutants from the zebrafish I. Dev Cell, 2004. 6(6): p. 771–80.

54. Pelegri, F., et al., Identification of recessive maternal-effect mutations in the zebrafish using a gynogenesis-based method. Dev Dyn, 2004. 231(2): p. 324–35.

55. Wagner, D.S., et al., Maternal control of development at the midblastula transition and beyond: mutants from the zebrafish II. Dev Cell, 2004. 6(6): p. 781–90.

56. Dosch, R., Next generation mothers: Maternal control of germline development in zebrafish. Crit Rev Biochem Mol Biol, 2015. 50(1): p. 54–68.

57. Hisano, Y., et al., Maternal and Zygotic Sphingosine Kinase 2 Are Indispensable for Cardiac Development in Zebrafish. J Biol Chem, 2015. 290(24): p. 14841–51.

58. Mendelson, K., et al., Maternal or zygotic sphingosine kinase is required to regulate zebrafish cardiogenesis. Dev Dyn, 2015. 244(8): p. 948–54.

59. Bassett, D.I., et al., Dystrophin is required for the formation of stable muscle attachments in the zebrafish embryo. Development, 2003. 130(23): p. 5851–60.

60. Hall, T.E., et al., The zebrafish candyfloss mutant implicates extracellular matrix adhesion failure in laminin alpha2-deficient congenital muscular dystrophy. Proc Natl Acad Sci U S A, 2007. 104(17): p. 7092–7.

61. Hall, T.E., et al., Cellular rescue in a zebrafish model of congenital muscular dystrophy type 1A. NPJ Regen Med, 2019. 4: p. 21.

62. Sun, Z., et al., Dyslipidemia in Muscular Dystrophy: A Systematic Review and Meta-Analysis. J Neuromuscul Dis, 2023. 10(4): p. 505–516.

63. Di Mauro, S., C. Trevisan, and A. Hays, Disorders of lipid metabolism in muscle. Muscle Nerve, 1980. 3(5): p. 369–88.

64. Saini-Chohan, H.K., et al., Delineating the role of alterations in lipid metabolism to the pathogenesis of inherited skeletal and cardiac muscle disorders: Thematic Review Series: Genetics of Human Lipid Diseases. J Lipid Res, 2012. 53(1): p. 4–27.

65. Amor, F., et al., Cholesterol metabolism is a potential therapeutic target in Duchenne muscular dystrophy. J Cachexia Sarcopenia Muscle, 2021. 12(3): p. 677–693.

66. Srivastava, N.K., et al., Abnormal lipid metabolism in skeletal muscle tissue of patients with muscular dystrophy: In vitro, high-resolution NMR spectroscopy based observation in early phase of the disease. Magn Reson Imaging, 2017. 38: p. 163–173.

67. Dabaj, I., et al., Muscle metabolic remodelling patterns in Duchenne muscular dystrophy revealed by ultra-high-resolution mass spectrometry imaging. Sci Rep, 2021. 11(1): p. 1906.

68. Vannoy, C.H., et al., Metabolomics Analysis of Skeletal Muscles from FKRP-Deficient Mice Indicates Improvement After Gene Replacement Therapy. Sci Rep, 2019. 9(1): p. 10070.

69. Sant, K.E. and A.R. Timme-Laragy, Zebrafish as a Model for Toxicological Perturbation of Yolk and Nutrition in the Early Embryo. Curr Environ Health Rep, 2018. 5(1): p. 125–133.

70. Miyares, R.L., V.B. de Rezende, and S.A. Farber, Zebrafish yolk lipid processing: a tractable tool for the study of vertebrate lipid transport and metabolism. Dis Model Mech, 2014. 7(7): p. 915–27.

71. Virote, B., et al., Obesity induction in adult zebrafish leads to negative reproduction and offspring effects. Reproduction, 2020. 160(6): p. 833–842.

72. Inoue, Y., et al., Maternal High-Fat Diet Affects the Contents of Eggs and Causes Abnormal Development in the Medaka Fish. Endocrinology, 2024. 165(3).

73. Chiara, V. and S.-Y. Kim, AnimalTA: A highly flexible and easy-to-use program for tracking and analysing animal movement in different environments. Methods in Ecology and Evolution, 2023. 14(7): p. 1699–1707.

74. Bailey, E.C., et al., NAD+ improves neuromuscular development in a zebrafish model of FKRP-associated dystroglycanopathy. Skelet Muscle, 2019. 9(1): p. 21.

75. Andrews, S. FastQC: A Quality Control Tool for High Throughput Sequence Data. 2010 [cited 2019; Available from: Https://Www.Bioinformatics.Babraham.Ac.Uk/Projects/Fastqc/.

76. Kim, D., B. Langmead, and S.L. Salzberg, HISAT: a fast spliced aligner with low memory requirements. Nat Methods, 2015. 12(4): p. 357–60.

77. Li, H., et al., The Sequence Alignment/Map format and SAMtools. Bioinformatics, 2009. 25(16): p. 2078–9.

78. Liao, Y., G.K. Smyth, and W. Shi, featureCounts: an efficient general purpose program for assigning sequence reads to genomic features. Bioinformatics, 2014. 30(7): p. 923–30.

79. Lawson, N.D., et al., An improved zebrafish transcriptome annotation for sensitive and comprehensive detection of cell type-specific genes. Elife, 2020. 9.

80. Love, M.I., W. Huber, and S. Anders, Moderated estimation of fold change and dispersion for RNA-seq data with DESeq2. Genome Biol, 2014. 15(12): p. 550.

81. Langfelder, P. and S. Horvath, WGCNA: an R package for weighted correlation network analysis. BMC Bioinformatics, 2008. 9: p. 559.

82. Zhang, B. and S. Horvath, A general framework for weighted gene co-expression network analysis. Stat Appl Genet Mol Biol, 2005. 4: p. Article17.

83. Shannon, P., et al., Cytoscape: a software environment for integrated models of biomolecular interaction networks. Genome Res, 2003. 13(11): p. 2498–504.

84. Lamprecht, M.R., D.M. Sabatini, and A.E. Carpenter, CellProfiler: free, versatile software for automated biological image analysis. Biotechniques, 2007. 42(1): p. 71–5.

