## Supplementary materials for "Impact of maternal compensation on developmental phenotypes in a zebrafish model of severe congenital muscular dystrophy"

Kyle P. Flannery et al.

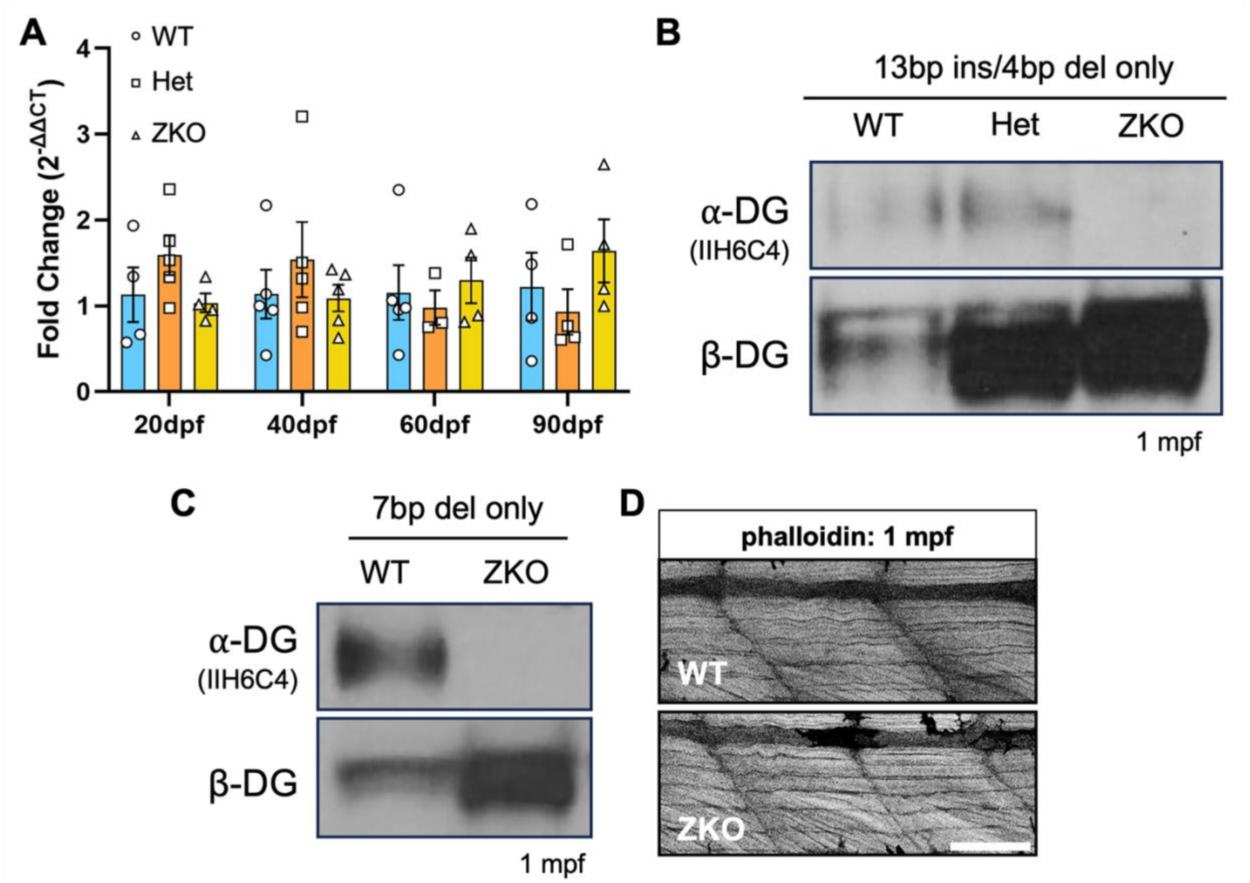


**Supplemental Figure 1**. ***Additional validation of the pomgnt2 line***. **A**: Quantitative real-time PCR analysis of *pomgnt2* gene expression showing no significant differences in WTs, Hets, and ZKOs. **B**-**C**: Western blot analysis of WGA enriched lysate showing that ZKOs with only the exon 1 (**B**) or exon 2 (**C**) mutations also show complete loss of α-DG glycosylation. **D**: Staining of muscle at 1 mpf with fluorescently conjugated phalloidin showing normal muscle fiber integrity in ZKOs (**Scale Bar**: 100 μm).

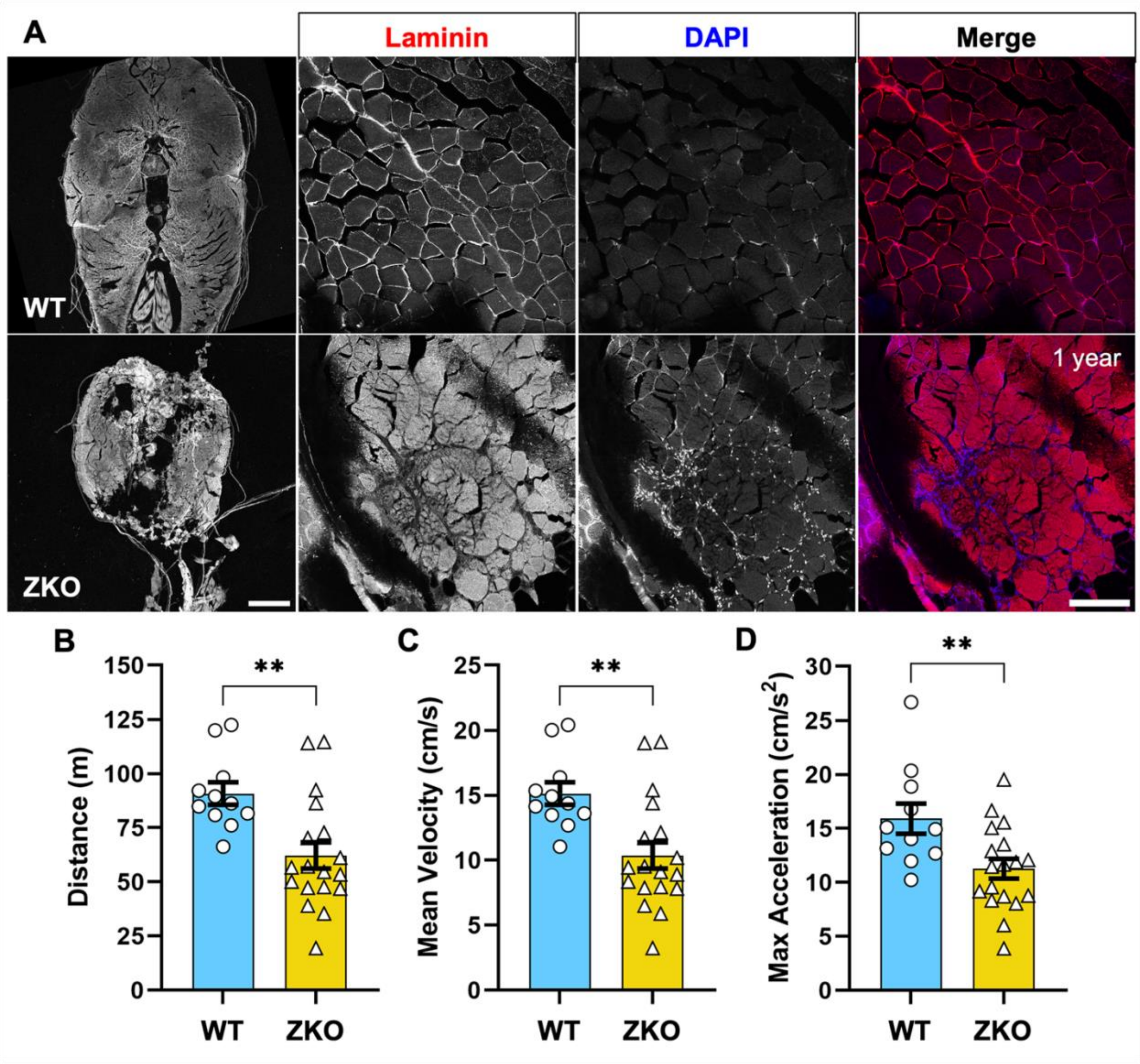


**Supplemental Figure 2**. **Muscle and motor phenotypes in 1 year old ZKOs. A**: Full transverse cryosections showing complete deterioration of muscle integrity in ZKOs reflective of advanced muscle disease, in addition to disrupted laminin staining around myofibers and increased nuclear staining (DAPI) suggestive of severe fibrosis (**Scale Bars**: 500 µm for whole cryosection, 20 µm for zoomed in images of muscle). **B-D**: Comprehensive analysis of swimming behavior and locomotor function showing that ZKOs have reductions in standard measures such as distance (**B**), velocity (**C**), and acceleration (**D**).

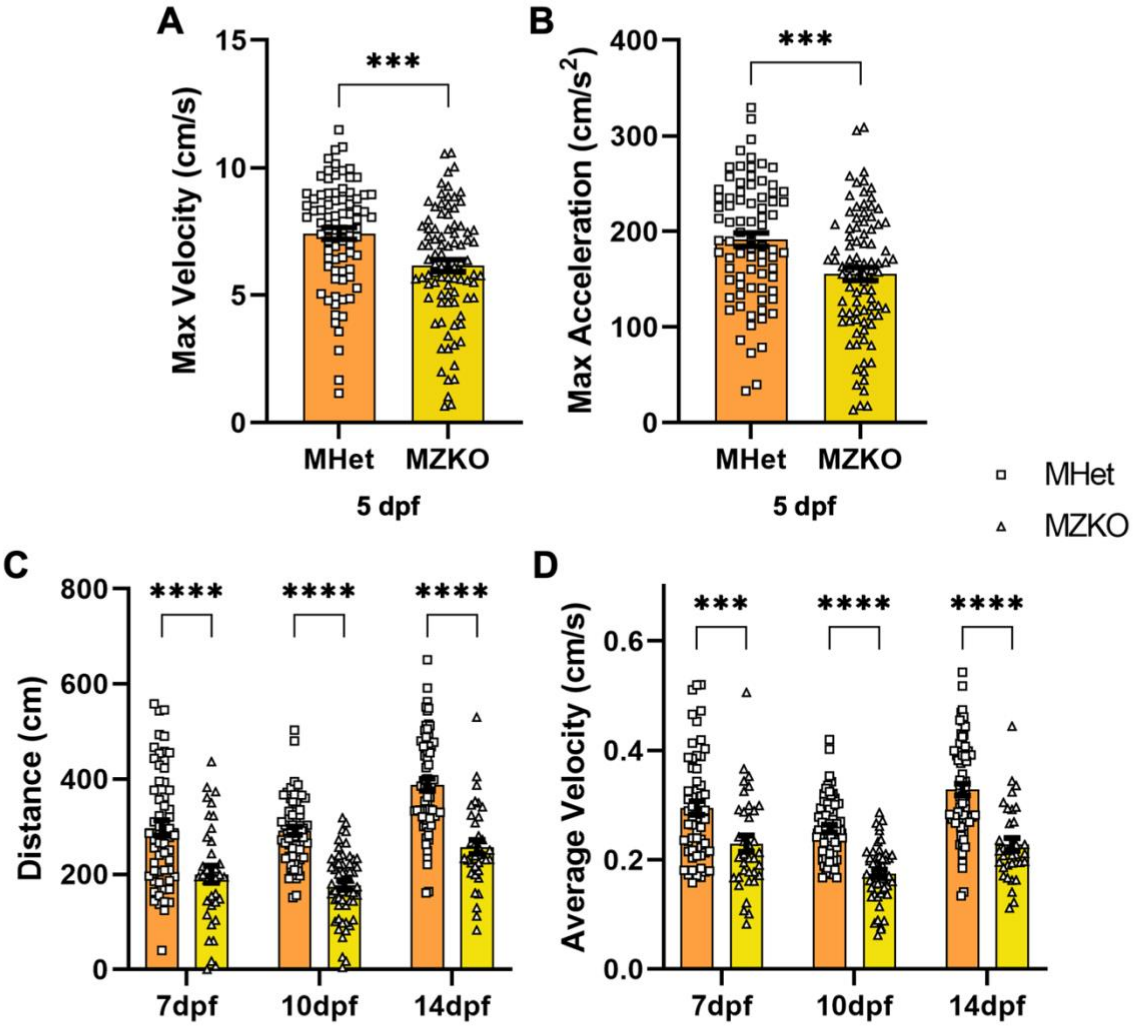


**Supplemental Figure 3**. **Additional assessment of locomotor function in MZKOs.**
**A**-**B**: Assessment of maximum velocity (**A**) and maximum acceleration (**B**) at 5 dpf showing significant reductions in the MZKOs. **C-D**: Assessment of total distance (**C**) and average velocity (**D**) showing significant reductions in the MZKOs at 7, 10, and 14 dpf.

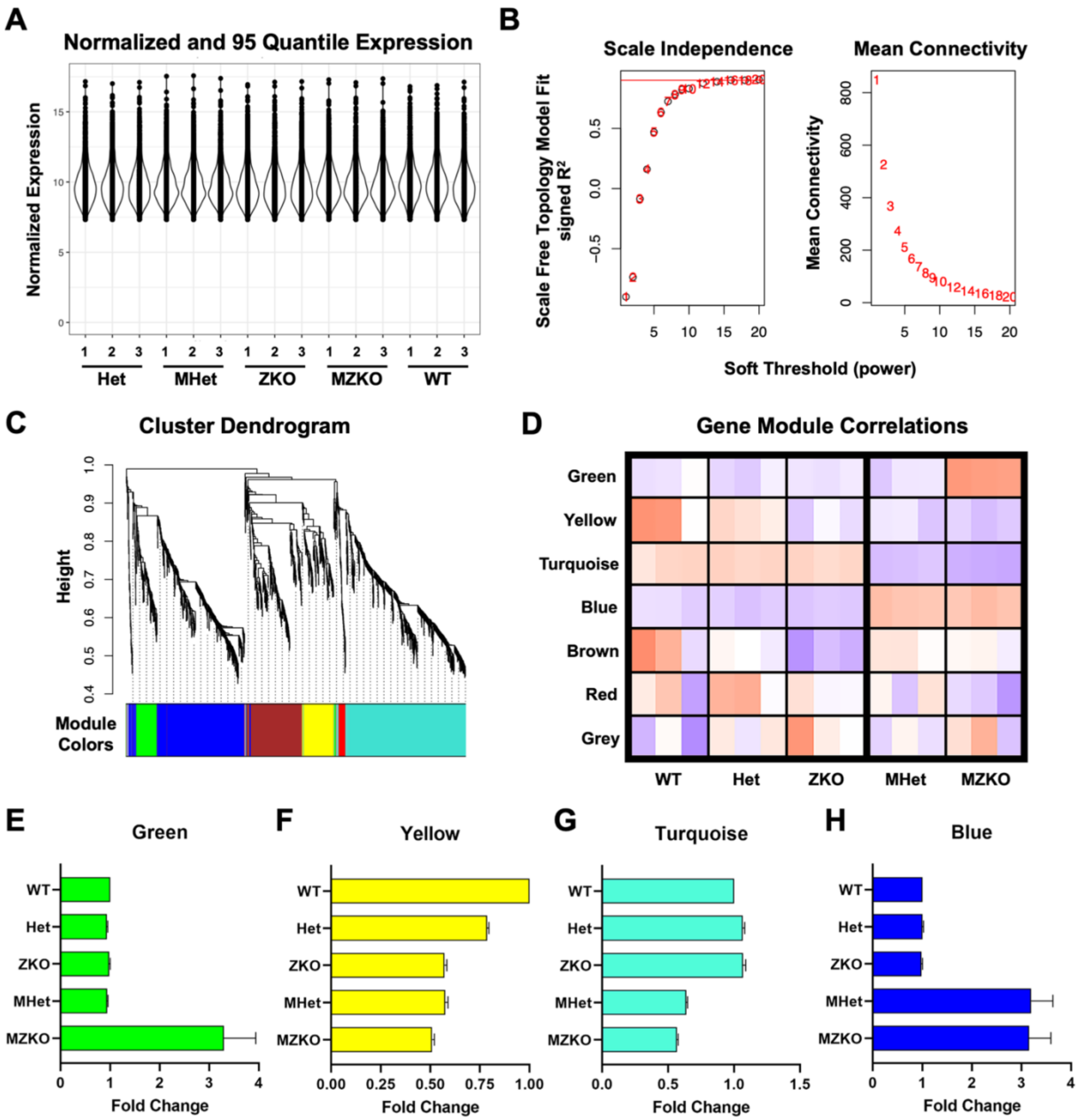


**Supplemental Figure 4**. ***Weighted gene co-expression network analysis pipeline and validation***. **A**: Data set normalization through DESeq2 and reduction to include genes expressed in the 95$^{th}$ quantile and above in all samples. **B**: Selection of soft thresholding power based on scale independence and mean connectivity. A soft thresholding value of 12 was selected. **C**: Dendrogram of gene modules derived from the reduced dataset of 1818 genes. The largest gene modules are turquoise and blue. **D**: Heat map of all gene modules identified, including those strongly correlated with maternal of zygotic genotype (green, yellow, turquoise, blue; also shown in **Fig. 3.9A**), and those that are not (brown, red, grey). **E-F**: Fold change of normalized expression in each module across genotypes compared to the WT samples as validation of gene module correlations.

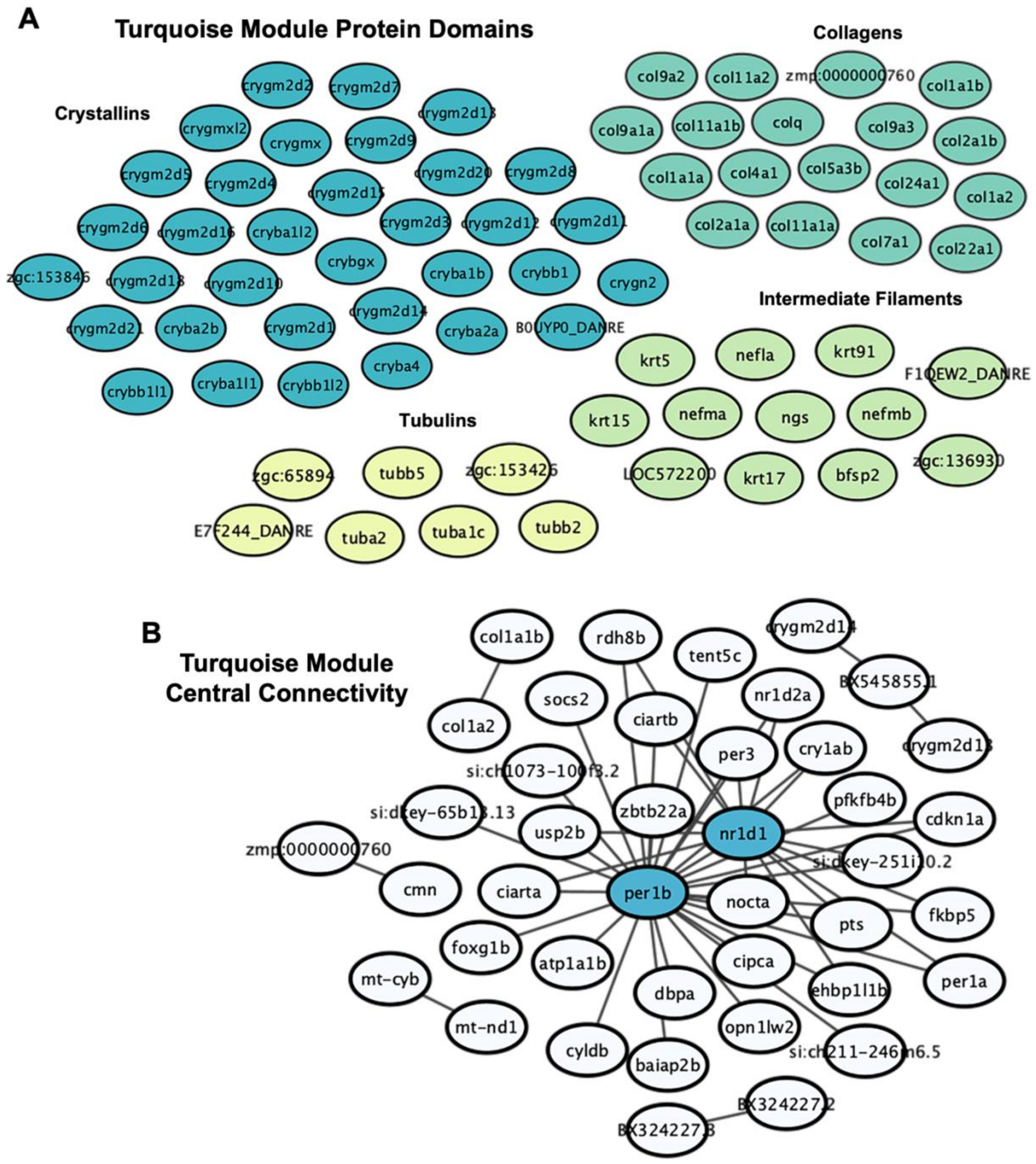


**Supplemental Figure 5.** ***Additional characterization of the turquoise module.*** **A**: Enrichment of crystallin, collagen, tubulin, and intermediate filament protein domains. **B**: Expression correlation analysis of the turquoise module revealing *per1b* and *nr1d1* to be the most centrally connected hub genes.

**Supplemental Table 1. Chi square analysis of survival in KOxHet crosses.**

| | ***Mpomgnt2* Het** | | ***MZpomgnt2* KO** | | | |
|---|---|---|---|---|---|---|
| **Timepoint** | **Observed** | **Expected** | **Observed** | **Expected** | $\chi^2$ | **p-value** |
| **10dpf** | 64 | 59 | 54 | 59 | 0.847 | 0.3573 |
| **14dpf** | 59 | 49 | 39 | 49 | 4.082 | *0.0434 |
| **28dpf** | 52 | 30.5 | 9 | 30.5 | 30.311 | ****<0.0001 |

**Supplemental Table 2.** Read mapping statistics for Chapter 3 RNA-sequencing experiments.

| Sample | Timepoint | Total Reads | Overall Alignment Rate (%) |
|---|---|---|---|
| WT1 | 5 dpf | 22221077 | 90.74 |
| WT2 | 5 dpf | 23382923 | 90.76 |
| WT3 | 5 dpf | 23294321 | 81.95 |
| Het1 | 5 dpf | 22365352 | 91.1 |
| Het2 | 5 dpf | 21146288 | 91.44 |
| Het3 | 5 dpf | 22645039 | 90.83 |
| ZKO1 | 5 dpf | 24740843 | 91.88 |
| ZKO2 | 5 dpf | 22103016 | 91.53 |
| ZKO3 | 5 dpf | 22659725 | 91.51 |
| MHet1 | 5 dpf | 21807697 | 91.6 |
| MHet2 | 5 dpf | 21035567 | 91.31 |
| MHet3 | 5 dpf | 21651629 | 87.64 |
| MZKO1 | 5 dpf | 20027272 | 91.13 |
| MZKO2 | 5 dpf | 22200195 | 91.04 |
| MZKO3 | 5 dpf | 22128356 | 91.06 |
| MHet1 | 10 dpf | 24497201 | 88.28 |
| MHet2 | 10 dpf | 27183778 | 87.01 |
| MHet3 | 10 dpf | 20628017 | 87.52 |
| MZKO1 | 10 dpf | 27251167 | 87.16 |
| MZKO2 | 10 dpf | 21329619 | 86.62 |
| MZKO3 | 10 dpf | 24302236 | 86.38 |
